# Temporal cortex astrocytic Gi-GPCR signaling regulates learned threat responses

**DOI:** 10.64898/2026.05.21.724789

**Authors:** Salome Nora Heimbach, Álvaro Collazos Matute, Victoria Steininger, Reuben Rajadhyaksha, Lea Klein, Lily Ferguson, Yaseer A. Sabir, Mingxia Huang, Alberto Cruz-Martín, Sarah Melzer

## Abstract

Astrocytes are increasingly recognized as dynamic modulators of brain circuit function, memory processing, and behavior. Emerging evidence suggests that astrocytic G-protein-coupled receptors (GPCRs) are key regulators of these processes through their influence on intracellular signaling and neuron-glia interactions.

Here, we show that the repertoire of functionally expressed GPCRs in cortical astrocytes is broader than previously appreciated. Yet, how distinct GPCR pathways contribute to behavioral regulation remains unknown for most brain areas and behavioral contexts. We therefore investigated the role of astrocytic GPCR signaling in the temporal cortex, a region that integrates multimodal sensory information and learned fear associations. Using chemogenetic tools to selectively activate distinct GPCR pathways in astrocytes, we demonstrate that Gi-coupled GPCR signaling, but not Gs- or Gq-coupled signaling, enhances fear memory retrieval. *In vivo* fiber photometry revealed that temporal cortex astrocytes exhibit robust Ca^2+^ transients to neutral, conditioned auditory and aversive sensory stimuli. Notably, astrocytic Gi-GPCR activation attenuated cue-evoked Ca^2+^ transients during memory retrieval.

Together, these findings identify astrocytic GPCR signaling as a pathway-specific regulator of fear memory retrieval and suggest that astrocytic Gi-GPCR signaling modulates the processing of sensory cues to drive defensive behavior.

## Main

Emotional memories, particularly fear memories, are an evolutionarily prioritized form of memory that support survival. Experiences associated with danger are acquired faster, stored robustly, and retrieved efficiently compared to neutral events (LeDoux, 1998; Mather et al., 2016; Tyng et al., 2017). Hence, classical fear conditioning paradigms in which memories of neutral cues can be acquired within a few trials through pairing with an aversive stimulus have become a key model to dissect how the brain encodes and retrieves emotional memories (LeDoux, 1998; Rogan et al., 1997).

The temporal association area (TeA), located in the temporal cortex ventral to auditory cortices and dorsal to ectorhinal and perirhinal cortices, is positioned to integrate auditory information with other sensory information and emotional signals. The TeA is connected with cortical regions spanning all sensory modalities, as well as subcortical regions that are commonly implicated in threat processing, including the amygdala and thalamus (Bedwell et al., 2015; Dalmay et al., 2019; Oh et al., 2014; Tasaka et al., 2020; Tovote et al., 2015; Zingg et al., 2014). Converging evidence indicates that the temporal cortex is required for the expression of auditory fear memories. Optogenetic inhibition of CaMKIIα^+^ temporal cortex neurons or their projections to the lateral amygdala during retrieval impairs fear expression (Dalmay et al., 2019). Similarly, chemogenetic suppression of vasoactive intestinal peptide (VIP)-expressing interneurons in the temporal cortex reduces fear expression (Cheng et al., 2025). Beyond its involvement in processing auditory information associated with aversive outcomes, the temporal cortex has been implicated in higher-order processing of sound concepts and of socially relevant sounds, including pup ultrasonic vocalizations that trigger maternal behavior (Tasaka et al., 2020; Trumpp et al., 2013). Together, these studies identify the temporal cortex as a higher-order cortical node that links auditory input to emotional valence.

Previous research on temporal cortex functions in fear memory has focused on the involvement of neuronal cell types. However, increasing evidence from other cortical as well as subcortical brain areas establishes astrocytes as key regulators of synaptic transmission, learning, memory, and synaptic plasticity (Adamsky et al., 2018; Araque et al., 2014; Savtchouk & Volterra, 2018). Several studies implicate astrocytic uptake of synaptic glutamate and potassium, and astrocytic release of gliotransmitters, most commonly glutamate, GABA, D-serine, and ATP, in these processes (Araque et al., 2014; Bergles & Jahr, 1997; Y. Li et al., 2020; Robin et al., 2018; Savtchouk & Volterra, 2018; Shen et al., 2022; Wu et al., 2022). Recent work further suggests that astrocytes can contribute to emotional memory in a region-specific manner, for example, by supporting fear-state-related neural representations in the amygdala (Bukalo et al., 2026).

Gliotransmission is evoked by elevations of Ca^2+^ in astrocytes (Araque et al., 2014). The dominant upstream regulators of astrocytic Ca^2+^ dynamics and their functional consequences vary across brain regions and behavioral contexts. *In vivo* studies in cortex reveal astrocytic GPCRs, including adrenergic receptors (Bekar et al., 2008; Oe et al., 2020; Paukert et al., 2014), metabotropic glutamate receptors (Wang et al., 2006), and histamine receptors (Taylor et al., 2025) as drivers of astrocytic Ca^2+^ elevations and gliotransmission. Additionally, evidence from subcortical areas or *in vitro* studies suggests that astrocyte function can be influenced by GPCRs for endocannabinoids (Navarrete & Araque, 2010), GABA (Shen et al., 2022; Yu et al., 2018), pituitary adenylate cyclase activating polypeptide (PACAP) (Kambe et al., 2021), and serotonin (González-Arias et al., 2023). However, for most of these GPCRs, the mechanisms that regulate astrocytic activity, gliotransmission, and function *in vivo* remain incompletely understood (Pereira et al., 2023).

Most astrocytic GPCRs fall into three major classes: Gq-, Gs-, and Gi-coupled receptors. Whereas Gq-and Gs-GPCRs are generally associated with increases in intracellular Ca^2+^ and cAMP levels, respectively, Gi-GPCR signaling was originally thought to reduce cAMP production. However, accumulating evidence indicates that activation of Gi-coupled GPCRs in astrocytes can also promote intracellular Ca^2+^ elevations, via mechanisms that are distinct from neuronal Gi signaling (Chai et al., 2017; Durkee et al., 2019; Nagai et al., 2019). Importantly, distinct astrocytic GPCR pathways can exert distinct functional effects (Vaidyanathan et al., 2021), highlighting the need for more rigorous dissection of GPCR signaling in astrocytes. These observations together suggest that astrocytic GPCR pathways serve as a critical interface through which modulators control neuronal activity, plasticity and memory processing (Drummond et al., 2024; Durkee et al., 2019; Y. Li et al., 2020; Shen et al., 2022).

Here, we combine *in vivo* and *in vitro* Ca^2+^ imaging and chemogenetic manipulation of astrocytes to investigate the role of GPCR signaling in temporal cortex astrocytes during auditory fear conditioning. We show that astrocytes exhibit robust Ca^2+^ elevations in response to conditioned auditory stimuli during fear retrieval, and that astrocytic Gi activation, but not Gs or Gq activation, enhances fear memory expression while modulating astrocyte dynamics in response to sensory cues. Together, these findings identify astrocytic GPCR signaling in the temporal cortex as a regulator of fear memory retrieval and suggest that distinct astrocytic GPCR pathways differentially shape how learned sensory cues influence defensive behavior.

## Results

### Astrocytes show widespread expression of GPCRs across multiple cortical areas and transcriptomic platforms

G-protein-coupled receptors (GPCRs) for glutamate and norepinephrine (NE) are well-known modulators of astrocyte function (Araque et al., 2014; Bekar et al., 2008). However, for the majority of GPCRs, expression patterns and functional relevance in astrocytes remain poorly defined.

To systematically assess GPCR expression in astrocytes, we analyzed publicly available single-cell RNA sequencing datasets (Sugino et al., 2019; Tasic et al., 2016; Yao et al., 2021), complemented by spatial transcriptomics (10 x Genomics Xenium Explorer 4.1.1) from sagittal mouse brain sections (Fig. 1a, Extended Data Fig. 1a). Astrocyte clusters were identified based on expression of canonical astrocytic markers (*Aqp4, Sox9, Aldh1l1*) and exclusion of neuronal (*Rbfox3*), ependymal (*Foxj1*), and perivascular fibroblast/oligodendrocyte progenitor cell (*Pdgfra*) markers (Fig. 1a, Extended Data Fig. 1b). We analyzed expression of 169 GPCRs present in most datasets (Fig. 1b, Extended Data Fig. 1b).

**Figure 1:**
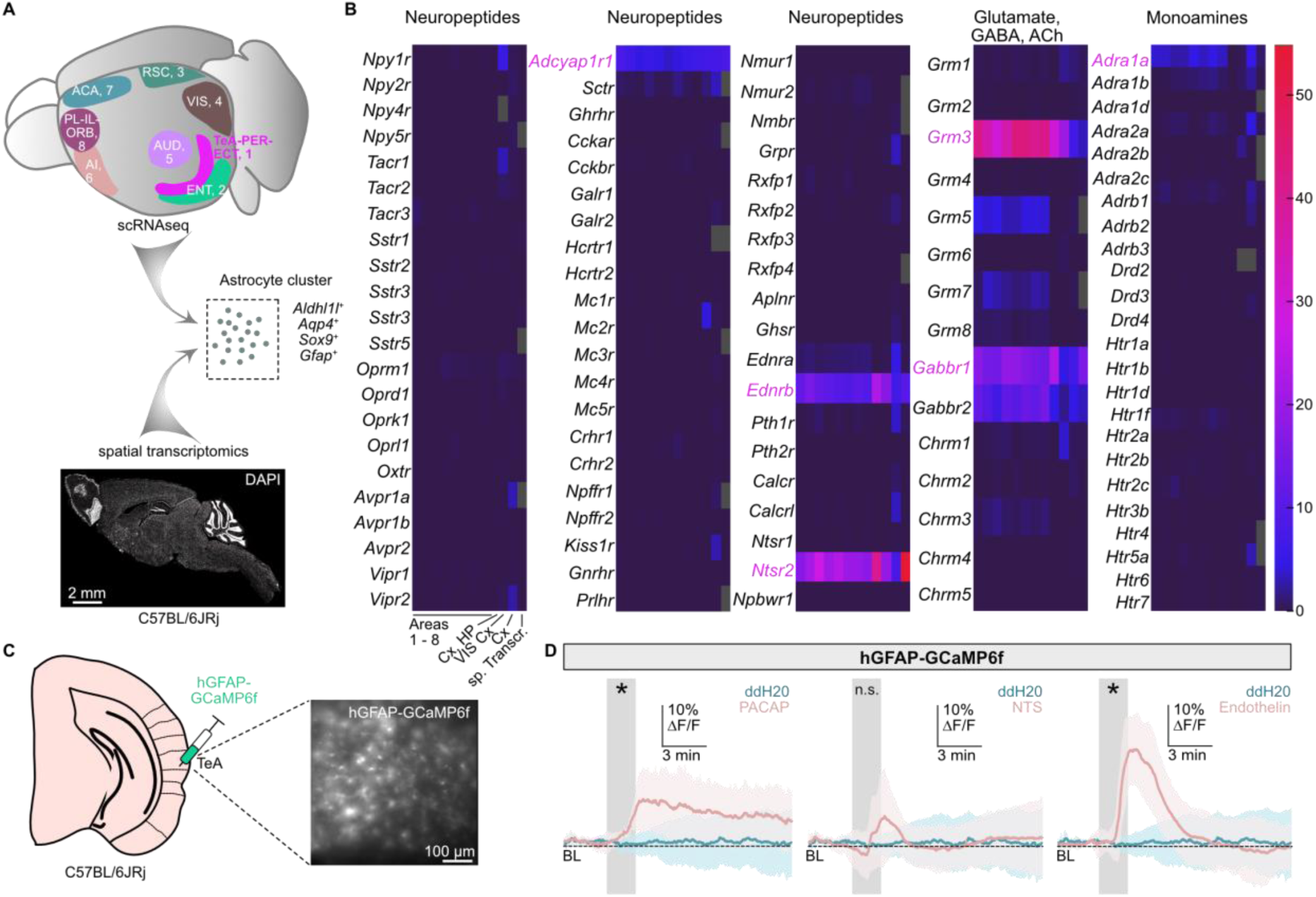
Astrocytes show GPCR expression across multiple cortical areas and transcriptomic platforms. (a) Schematic of workflow for GPCR expression analysis in astrocytic clusters. Cortical areas that were analyzed, and visualized in Fig. 1b, are highlighted. (b) Heatmaps showing normalized expression of selected GPCR genes in astrocyte clusters across transcriptomic datasets. Columns 1-8 correspond to cortical regions highlighted in Fig. 1a, data originally published in Yao et al. (2021). Columns 9-11 show averaged expression in indicated brain areas. Data from Yao et al. (2021), Tasic et al. (2016), and Sugino et al. (2019). Column 12 represents astrocyte-rich clusters identified in spatial transcriptomic data from sagittal mouse brain sections. Color scale indicates normalized expression within each dataset. Grey fields indicate data not available. (c) Schematic of TeA astrocytic Ca^2+^ imaging experiment. Exemplary epifluorescence image of GCaMP6f labeling in an acute TeA slice. (d) Astrocytic Ca²⁺ imaging in acute TeA slices expressing AAV-hGFAP-GCaMP6f. Ligand application is indicated by the grey shaded area. Peak signals (mean ± SEM): pituitary adenylate cyclase activating polypeptide (PACAP) 13.68% ± 1.18% ΔF/F; neurotensin (NTS) 9.72% ± 1.11% ΔF/F; endothelin 25.76% ± 1.42% ΔF/F; ddH_2_O 3.4% ± 0.59%. Significant increases in GCaMP6f fluorescence indicated by asterisks (PACAP: *p* = 0.043; NTS: *p* = 0.073; endothelin: *p* ≤ 0.01; unpaired *t*-test, *N* = 6 slices per condition). Abbreviations: ACA, anterior cingulate cortex; AI, anterior insula; AUD, auditory cortex; Cx, Cortex; ENT, entorhinal cortex; HP, hippocampus; PL-IL-ORB, prelimbic, infralimbic, orbitofrontal cortex; RSC, retrosplenial cortex; TeA-PER-ECT, temporal association, perirhinal, ectorhinal cortex; VIS, visual cortex. Data are presented as mean ± SEM.

The most prominently expressed GPCRs were the neuropeptide receptors *Ntsr2, Ednrb,* and *Adcyap1r1*, the metabotropic glutamate receptor *Grm3,* the GABA-B receptor *Gabbr1*, the norepinephrine receptor *Adra1a*, and the lipid receptor *S1pr1*. These GPCRs were highly expressed across multiple cortical regions and were also detected in the spatial transcriptomics dataset, which included cells from the cortex, hippocampus, amygdala, striatum, thalamus, midbrain, and cerebellum. This suggests that the identified GPCR profile is not restricted to a single anatomical region. Notably, among the most highly expressed GPCRs, receptors predicted to couple to Gi/o proteins, including *Ntsr2*, *Ednrb*, *Grm3*, *Gabbr1*, and *S1pr1* (Bettler et al., 2004; Horstmeyer et al., 1996; Pin & Duvoisin, 1995; Rosen et al., 2013; Vita et al., 1998), were overrepresented relative to receptors predicted to couple to Gq- or Gs proteins (Inoue et al., 2019). Additional analysis revealed that astrocytes showed strong expression of *Gnao1*, encoding the Gi/o-family Gαo subunit, suggesting that astrocytes possess the molecular machinery required for Gi/o-mediated signal transduction (Extended Data Fig. 1b). Together, these data establish a previously underappreciated GPCR landscape in the cortex and provide the rationale for testing whether astrocyte GPCR signaling supports cortex function.

To test whether these transcriptionally identified GPCRs are functionally expressed in astrocytes, we examined the TeA, a higher-order cortical region involved in integrating sensory input and emotional valence (Cheng et al., 2025; Dalmay et al., 2019; Tasaka et al., 2020). Therefore, we performed Ca^2+^ imaging in acute TeA brain slices expressing AAV-hGFAP-GCaMP6f (Fig. 1c) while blocking neuronal activity. In previous studies, Ca^2+^ elevations were frequently observed downstream of astrocytic Gi-and Gq-GPCR signaling (Durkee et al., 2019; Wang et al., 2006). Bath application of ligands for highly expressed receptors – including neurotensin (NTS; *Ntsr2*), endothelin (*Ednrb*), glutamate (*Grm3*), pituitary adenylate cyclase-activating polypeptide (PACAP; *Adcyap1r1*), and norepinephrine (*Adra1a*) – resulted in intracellular Ca^2+^ elevations in astrocytes (Fig. 1d, Extended Data Fig. 1c). These findings indicate that GPCRs identified at the transcriptional level can engage intracellular signaling pathways in cortical astrocytes.

Together, these results demonstrate that diverse neuropeptidergic and neuromodulatory GPCRs can drive intracellular Ca^2+^ signaling in cortical astrocytes.

### Astrocytic Gi-GPCR activation in TeA enhances fear memory retrieval

Previous work has established that the temporal neocortex is required for the expression of learned auditory fear memories, particularly in tasks where mice are subjected not only to conditioned sounds but also to ‘distractor’ sounds that are not paired with a shock (Dalmay et al., 2019). Neurons in the temporal neocortex preferentially encode higher-order acoustic features and show enhanced engagement during auditory discrimination and associative learning involving frequency-modulated stimuli and complex sound patterns (Dalmay et al., 2019; Feigin et al., 2021). Based on these findings, we employed an auditory fear conditioning paradigm using upward and downward frequency sweeps as conditioned (CS^+^) and ‘distractor’ (CS^-^) stimuli to test the function of GPCR signaling in cortical astrocytes. After one day of habituation, mice were subjected to 15 CS^+^ co-terminating with mild foot shocks and 15 CS^-^. Fear memory retrieval was tested 24 h later (Fig. 2a).

**Figure 2:**
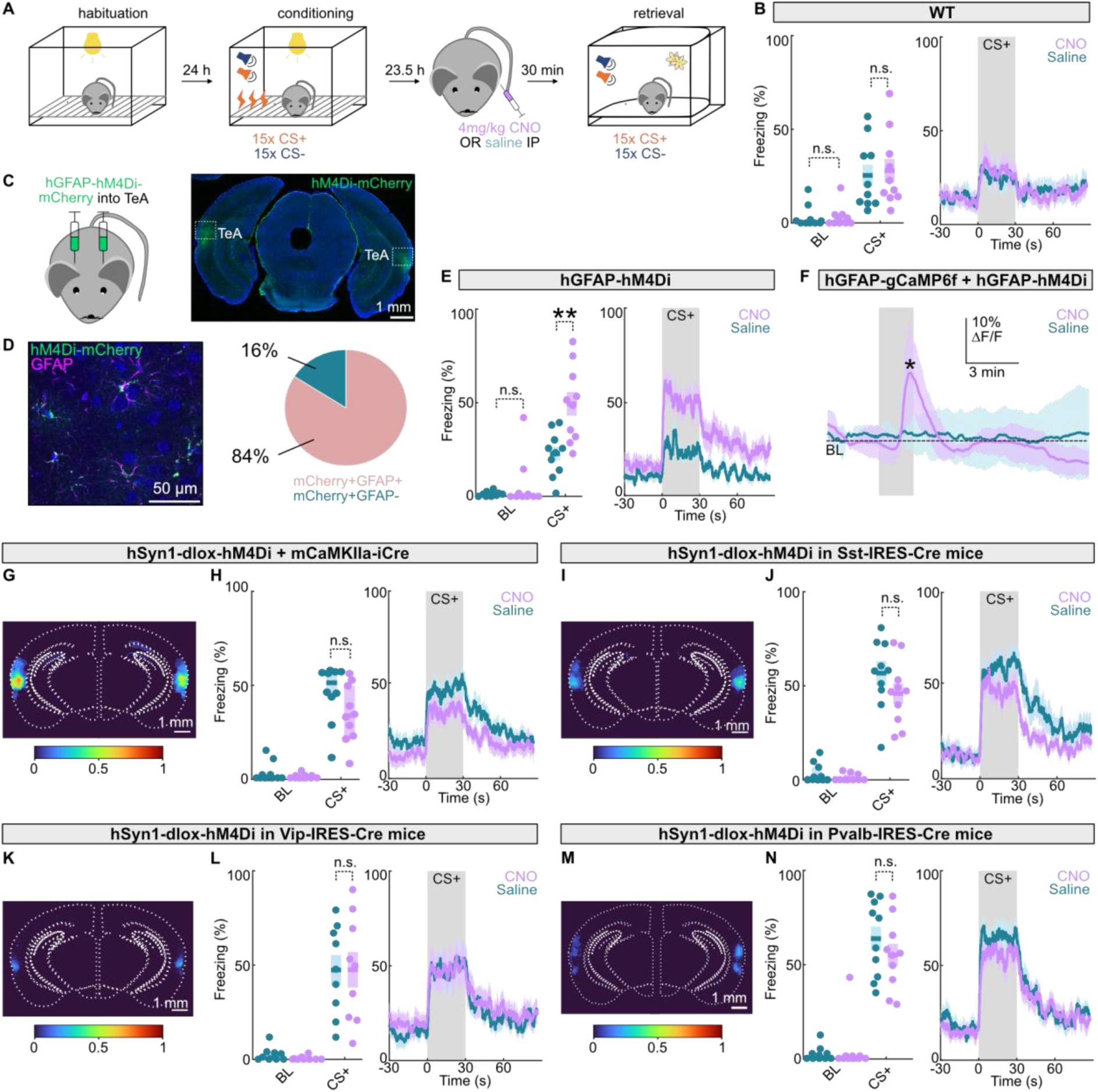
Astrocytic Gi-GPCR activation in the TeA enhances fear memory retrieval. (a) Schematic of discriminatory auditory fear conditioning paradigm with paired (CS⁺) and unpaired (CS^-^) auditory sweeps. CNO or saline was administered 30 min prior to memory retrieval testing. (b) Control experiment to test off-target CNO effects in naïve wild-type mice. Left: baseline freezing (median [IQR]; CNO: 0.87% [0.00-4.40%], saline: 0.56% [0.00-1.74%]; *p* = 1, Mann-Whitney *U* test) and CS⁺ freezing (mean ± SEM; CNO: 27.93 ± 6.25%, saline: 25.39 ± 5.74%; *p* = 1, unpaired *t-*test) for saline-and CNO-treated animals. Each dot represents one mouse. Right: Peri-stimulus freezing time course aligned to CS⁺ onset (CS⁺ interval shaded), *N* = 10 mice per condition. (c) Schematic and exemplary epifluorescence image showing injection of AAV-hGFAP-hM4D_i_-mCherry into TeA. (d) Exemplary confocal image showing immunostaining with GFAP to validate co-expression. Quantification is shown on the right. *N* = 6 slices from 3 mice. (e) Freezing during retrieval following astrocytic Gi-GPCR activation. Left: Freezing quantification during baseline (CNO: 0.00% [0.00-1.65%], saline: 1.16% [0.00-1.74%]; *p* = 0.842; Mann-Whitney *U* test) and CS⁺ for saline- and CNO-treated groups (CNO: 49.41 ± 6.36%; saline: 22.78% ± 3.77%; *p* ≤ 0.01; unpaired *t*-test). Right: freezing time course aligned to CS⁺ onset, *N* = 10 per condition. (f) Astrocytic Ca²⁺ imaging in acute TeA slices co-expressing AAV-hGFAP-GCaMP6f and hGFAP-hM4D_i_-mCherry. CNO or saline application is indicated by the grey shaded area. Mean peak signals: CNO 13.53 ± 1.33% ΔF/F, saline 3.55 ± 0.75% ΔF/F. CNO significantly increased GCaMP6f fluorescence (*p* = 0.0234; unpaired *t*-test), *N* = 6 slices per condition. (g) Overlay of viral injection sites for AAV-hSyn1-dlox-hM4D_i_-mCherry and mCaMKIIα-iCre in TeA, for mice used in behavioral tests. Color scale indicates proportion of mice with detectable virus expression. (h) Freezing during retrieval following chemogenetic Gi-GPCR activation of TeA CaMKII^+^ neurons. Left: Quantification of freezing during baseline and CS⁺ (CNO: 33.76% [22.96-49.69%], saline: 51.31% [43.77-57.12%]; *p* = 0.162, Mann-Whitney *U* test). Right: CS⁺-aligned freezing time course for saline-and CNO-treated groups, *N* = 10 mice per condition. (i) Same as in (g), but for AAV-hSyn1-dlox-hM4D_i_-mCherry expression in Sst-IRES-Cre mice. (j) Same as in (h), but for *Sst⁺* neurons (CS^+^ CNO: 45.88% ± 1.72%; saline: 55.83% ± 1.84%, *p* = 0.640, unpaired *t*-test), *N* = 10 per condition. (k) Same as in (g), but for AAV-hSyn1-dlox-hM4D_i_-mCherry expression in Vip-IRES-Cre mice. (l) Same as in (h), but for *Vip⁺* neurons (CS^+^ CNO: 47.51% ± 3.14%; saline: 47.40% ± 2.64%, *p* = 0.993, unpaired *t*-test), *N* = 9 per condition. (m) Same as in (g), but for AAV-hSyn1-dlox-hM4D_i_-mCherry in Pvalb-IRES-Cre mice. (n) Same as in (h), but for *Pvalb⁺* neurons (CS^+^ CNO: 54.91% ± 5.93%; saline: 63.64% ± 6.41%, *p* = 0.992, unpaired *t*-test), *N* = 10 per condition.

To specifically activate Gi-GPCR signaling, we made use of the designer receptor hM4D_i_ and its ligand clozapine N-oxide (CNO; 4 mg/kg i.p.). We excluded nonspecific effects of CNO on fear memory retrieval by examining the effect of systemic CNO in naïve control mice. CNO had no detectable effect on baseline freezing levels, or freezing levels in response to the CS⁺ and CS^-^, or the discrimination index during the retrieval session (Fig. 2b, Extended Data Fig. 2a,b), indicating that the behavioral paradigm is insensitive to putative off-target effects of CNO. Freezing to CS^+^ (but not CS^-^) is dependent on temporal cortex function (Dalmay et al., 2019). Therefore, all subsequent analyses focused on CS^+^. Because most experiments were performed in female mice and the estrous stage can introduce behavioral variability, we assessed fear memory retrieval across the estrous cycle. Freezing to the CS^+^ during retrieval did not differ across estrous stages (Extended Data Fig. 2c).

To selectively activate Gi-GPCR signaling in astrocytes, we injected AAV-hGFAP-hM4D_i_-mCherry into the temporal neocortex, focusing on the TeA (hereafter called TeA; Fig. 2c). Immunohistochemical analysis confirmed that 84% of virally labeled cells were GFAP^+^ astrocytes (Fig. 2d). Chemogenetic activation of astrocytic Gi-GPCR signaling significantly increased freezing to the CS^+^ during retrieval (*p* ≤ 0.01, Fig. 2e; Extended Data Fig. 2d,e). This effect was not explained by differences in learning. Thus, CNO- and control groups acquired the tone-shock association similarly during conditioning, as indicated by comparable levels of freezing during the last CS^+^ presentation (Extended Data Fig. 2f). Moreover, both groups exhibited similar shock responses during conditioning (*p* = 0.8; Extended Data Fig. 2g), indicating equivalent shock sensitivity prior to hM4D_i_ activation.

We next tested whether hM4D_i_ activation alters astrocyte Ca²⁺ dynamics by co-expressing AAV-hGFAP-GCaMP6f and imaging Ca²⁺ signals in acute TeA slices. Bath application of 10 µM CNO produced a transient but significant increase in astrocytic Ca²⁺ (*p* = 0.023; Fig. 2f).

To determine whether the behavioral effect was specific to astrocytic Gi-GPCR signaling, we chemogenetically activated Gi-GPCRs in multiple neuronal populations. To target excitatory neurons, we co-injected AAV-hSyn1-dlox-hM4D_i_-mCherry and AAV-mCaMKIIα-iCre into the TeA of wild-type mice (Fig. 2g). Chemogenetic activation of hM4D_i_ in CaMKIIα^+^ neurons did not alter CS^+^ freezing during retrieval (Fig. 2h, Extended Data Fig. 2h).

We next examined major classes of inhibitory interneurons by injecting AAV-hSyn1-dlox-hM4D_i_-mCherry into Sst-IRES-Cre, Vip-IRES-Cre, and Pvalb-IRES-Cre mice (Fig. 2i,k,m). Since *Pvalb*^+^ interneurons are sparse in TeA (Cheng et al., 2025), viral expression in Pvalb-IRES-Cre mice was observed mainly dorsal and ventral to the TeA (Fig. 2m). Gi-GPCR activation in *Sst*^+^ cells (Fig. 2j, Extended Data Fig. 2i), *Vip*^+^ cells (Fig. 2l, Extended Data Fig. 2j), or *Pvalb*^+^ cells (Fig. 2n, Extended Data Fig. 2k) did not significantly affect CS⁺ freezing.

As these experiments were performed with in-house bred mouse lines, we repeated the CNO control experiments in non-transgenic naïve littermates. Again, we did not detect any off-target effects of CNO, as indicated by comparable CS^+^ freezing between groups during the retrieval session (Extended Data Fig. 2l).

Together, these results identify astrocytic Gi-GPCR signaling in the TeA as a central regulator of fear memory retrieval.

### Gq- and Gs-GPCR activation have no significant effect on fear memory retrieval

Since our data suggested expression of putative Gs- and Gq-GPCRs in temporal cortex astrocytes (Fig. 1), we next tested whether activating astrocytic Gq- or Gs-GPCR signaling in TeA is sufficient to modulate fear memory retrieval using the same fear conditioning paradigm as above. To activate Gq-GPCR signaling, we expressed AAV-hGFAP-hM3D_q_-mCherry in TeA astrocytes (Fig. 3a). During retrieval, CNO administration did not significantly alter freezing to the CS^+^ compared with saline-treated controls (Fig. 3b, Extended Data Fig. 3a), although a trend in the same direction as for Gi-GPCR activation was observed.

**Figure 3:**
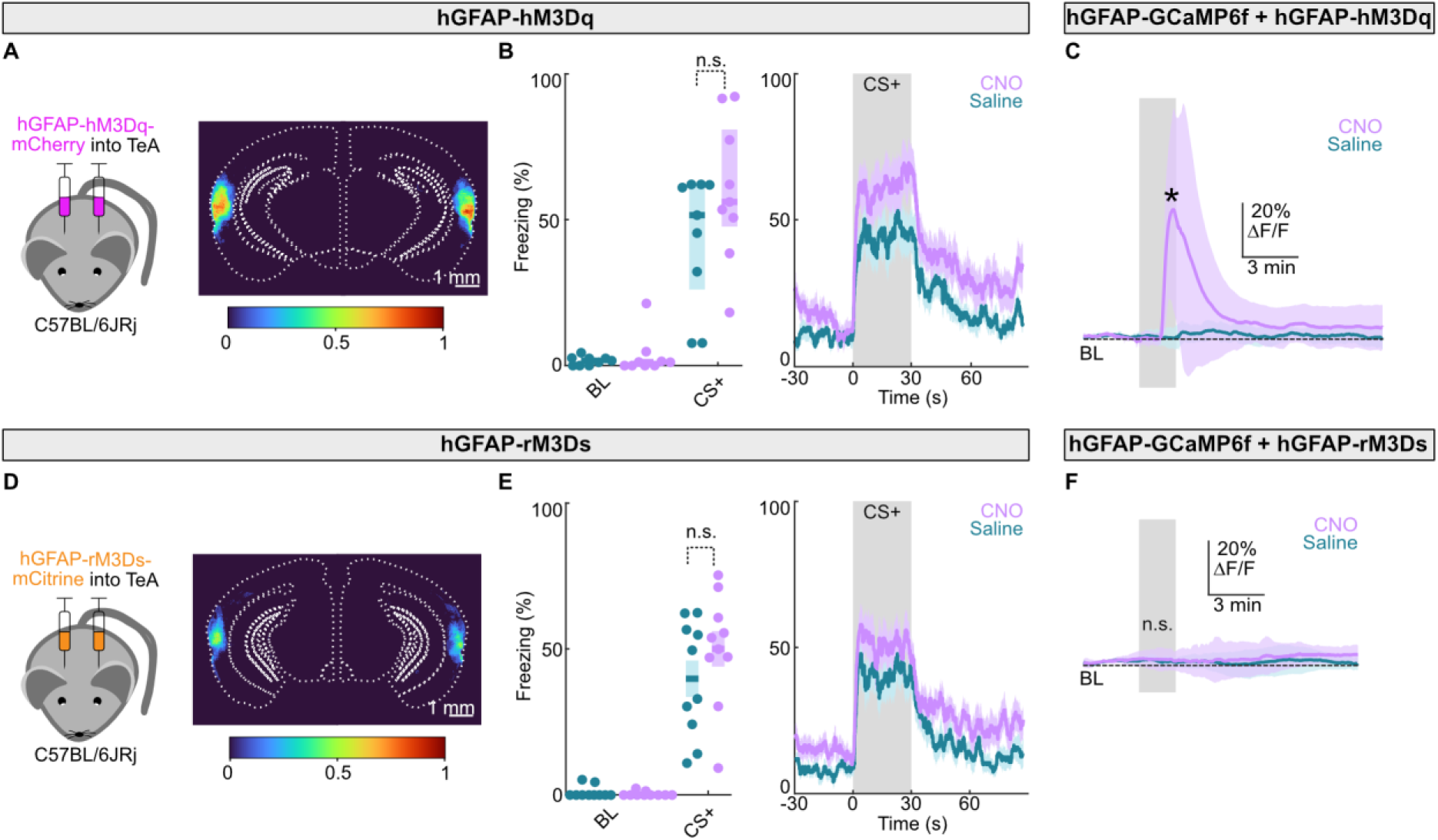
Gq- and Gs-GPCR activation have no significant effect on fear memory retrieval. (a) Schematic and overlay of main viral injection sites for AAV-hGFAP-hM3D_q_-mCherry in TeA, for mice used in behavioral tests. Color scale indicates proportion of mice with detectable virus expression. (b) Freezing during retrieval following astrocytic Gq-GPCR activation. Left: freezing quantification during baseline and CS⁺ for saline- and CNO-treated groups (CS^+^: CNO: 56.08% [47.64-80.92%]; saline: 51.56% [26.11-61.98%], *p* = 0.867, Mann-Whitney *U* test). Right: freezing time course aligned to CS⁺ onset, *N* = 9 mice per condition. (c) Astrocytic Ca²⁺ dynamics in acute TeA slices co-expressing AAV-hGFAP-GCaMP6f and AAV-hGFAP-hM3D_q_-mCherry. CNO/saline application is indicated by the grey shaded area. Peak signal: CNO: 47.43 ± 6.72% ΔF/F, saline: 3.54 ± 0.61% ΔF/F. CNO significantly increased GCaMP6f fluorescence (*p* = 0.024, unpaired *t*-test), *N* = 6 slices per condition. (d) Schematic and overlay of main viral injection sites for AAV-hGFAP-HA-rM3D_s_-mCitrine in TeA, for mice used in behavioral tests. Color scale indicates proportion of mice with detectable virus expression. (e) Same as in (b), but for mice expressing AAV-hGFAP-HA-rM3D_s_-mCitrine (CS^+^ CNO: 50.17% ± 6.08%; saline: 39.90% ± 6.25%, *p* = 0.763, unpaired *t*-test), *N* = 10 mice per condition. (f) Astrocytic Ca²⁺ dynamics in acute TeA slices co-expressing AAV-hGFAP-GCaMP6f and AAV-hGFAP-HA-rM3D_s_-mCitrine, peak signal: CNO: 3.15% [2.01-6.99%] ΔF/F, saline: 2.78% [0.82-2.87%] ΔF/F (*p* = 0.738; Mann-Whitney *U* test), *N* = 6 slices per condition.

*In vitro*, astrocytic Gq-GPCR activation produced robust Ca²⁺ elevations in acute TeA slices co-expressing AAV-hGFAP-GCaMP6f and AAV-hGFAP-hM3D_q_-mCherry (*p* = 0.024; Fig. 3c), confirming the functionality of the tool. We next examined astrocytic Gs-GPCR signaling by expressing AAV-hGFAP-HA-rM3D_s_-mCitrine in TeA astrocytes (Fig. 3d). Similar to Gq-GPCR activation, Gs-GPCR activation showed a trend toward increased CS^+^-evoked freezing during retrieval, but this effect did not reach statistical significance (Fig. 3e, Extended Data Fig. 3b). In acute TeA slices, rM3D_s_ activation did not produce increased Ca²⁺ transients (Fig. 3f) as expected for Gs-GPCR signaling (Oe et al., 2020).

These results indicate that astrocytic Gi-GPCR activation robustly enhances fear retrieval, whereas Gq-and Gs-GPCR activation have minor, non-significant effects, suggesting that different GPCR pathways differentially affect fear memory retrieval.

### TeA astrocytes exhibit sensory-evoked Ca2+ dynamics that lag neuronal activity

To directly visualize Ca^2+^ dynamics *in vivo*, we used fiber photometric imaging of TeA astrocytes during auditory fear conditioning and retrieval (Fig. 4a). To this end, AAV-hGFAP-GCaMP6f was injected into the TeA of wild-type mice (Fig. 4b). During fear conditioning, GCaMP6f signals increased in response to both CS^+^ and CS^−^ (Fig. 4c), with the largest amplitudes following foot shock delivery (Fig. 4c). CS^+^ responses showed no significant attenuation or amplification across the 15 conditioning trials (*p* = 0.866; Extended Data Fig. 4a). During retrieval, astrocytes exhibited stimulus-evoked Ca²⁺ transients to both CS^+^ and CS^-^ (Fig. 4d), with no significant attenuation across trials (*p* = 0.237; Extended Data Fig. 4b).

**Figure 4:**
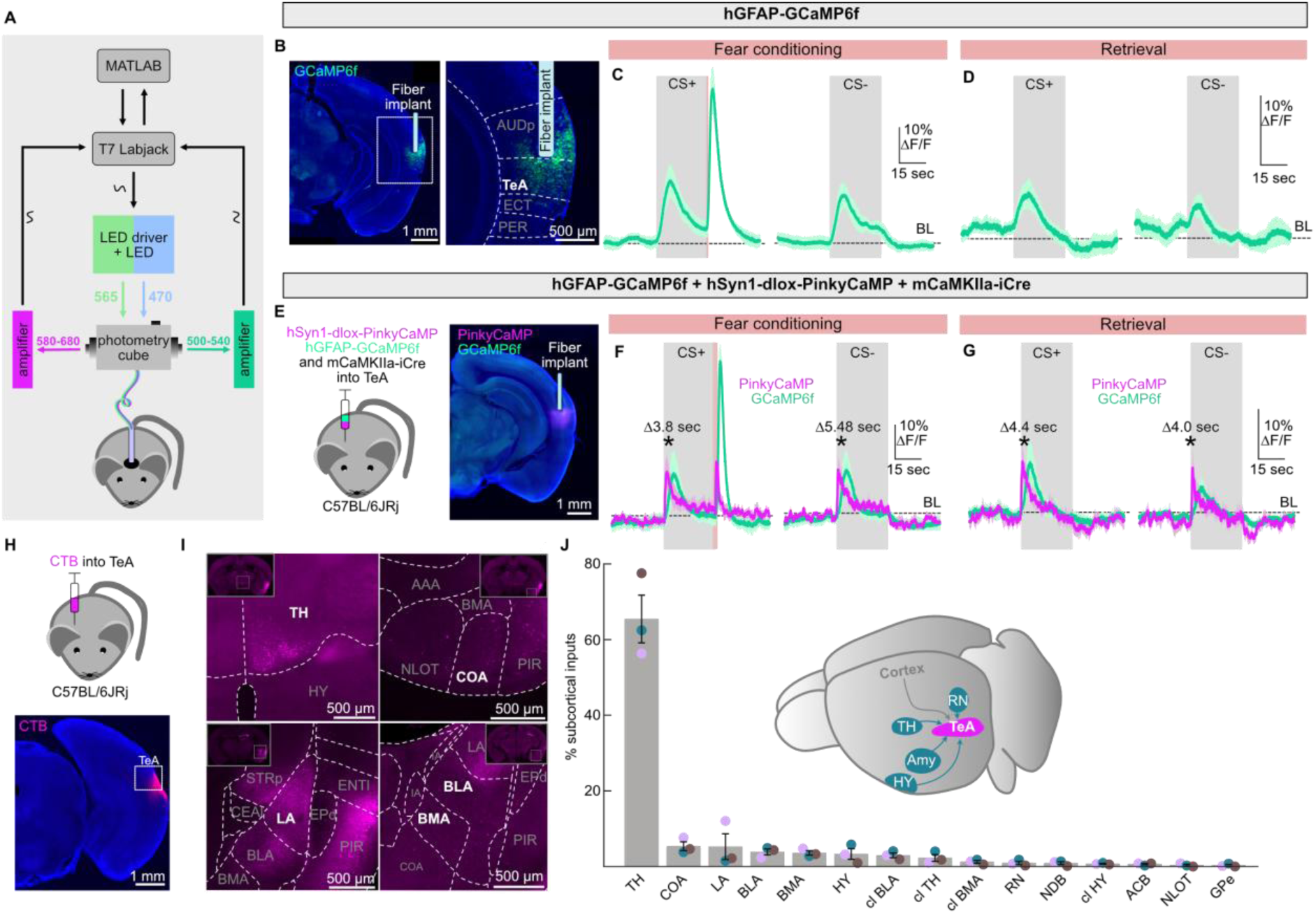
T**e**A **astrocytes exhibit sensory-evoked Ca^2+^ dynamics that lag neuronal activity**. (a) Schematic of fiber photometry setup used to record Ca^2+^ signals in the TeA *in vivo*. (b) Exemplary epifluorescence image showing AAV-hGFAP-GCaMP6f expression and fiber implant location. (c) Normalized fluorescence changes (ΔF/F) from AAV-hGFAP-GCaMP6f-expressing astrocytes aligned to CS⁺ (peak ΔF/F: 18.2 ± 1.27%) and CS^−^ (peak ΔF/F: 14.54 ± 0.78%) presentations during fear conditioning. The sounds and foot shock are indicated by grey and peach-colored vertical bars (peak ΔF/F after shocks: 42.81 ± 2.67%), *N* = 6 mice. (d) Normalized fluorescence changes (ΔF/F) from AAV-hGFAP-GCaMP6f-expressing astrocytes aligned to CS⁺ and CS^−^ presentations during retrieval (peak ΔF/F CS^+^: 6.26 ± 0.58%; peak ΔF/F CS^-^: 4.52 ± 0.56%), *N* = 6 mice. (e) Schematic and representative epifluorescence image showing AAV-hGFAP-GCaMP6f and AAV-hSyn1-dlox-PinkyCaMP expression, and fiber implant location in TeA. (f) Normalized fluorescence changes (ΔF/F) from AAV-hGFAP-GCaMP6f-expressing astrocytes and AAV-hSyn1-dlox-PinkyCaMP-expressing neurons aligned to CS⁺ (left) and CS^−^ (right) presentations during fear conditioning. Peak ΔF/F for astrocytes during CS^+^: 12.82 ± 3.00%; latency: 5.2 ± 0.17 s; peak ΔF/F for neurons: 13.25 ± 2.06%; latency: 1.4 ± 0.09 s. Peak ΔF/F for astrocytes during CS^-^: 16.63 ± 4.81%; latency: 6.83 ± 0.63 s; peak ΔF/F for neurons: 15.17 ± 3.60%; latency: 1.35 ± 0.39 s. Foot shock (peach bar) evoked large astrocytic (46.5 ± 2.28%) and neuronal (16.27 ± 0.80%) responses. Peak latency differed significantly between astrocytes and neurons for both CS^+^ and CS^-^ (both *p* ≤ 0.01; paired *t*-test), *N* = 5 mice. (g) Same as (f), but for fear retrieval. Peak ΔF/F for astrocytes during CS^+^: 10.11 ± 0.27%; latency 5.3 ± 0.27 s; peak ΔF/F for neurons: 10.28 ± 1.35%; latency 0.9 ± 0.27 s. CS^-^: Peak ΔF/F astrocytes: 3.42% [2.97-7.95%]; latency 4.78 ± 0.1 s. Peak ΔF/F neurons 10.74% [6.87-15.84%]; latency 0.78 ± 0.11 s. Peak latency differed significantly between astrocytes and neurons for CS^+^ and CS^-^ (both *p* ≤ 0.01; paired *t*-test), *N* = 5 mice. (h) Schematic and exemplary injection site of CTB in TeA. (i) Exemplary epifluorescence images of CTB signal in TH, COA, LA, BLA, and BMA. (j) Schematic and quantification of the proportion of CTB^+^ cell bodies detected in the analyzed subcortical areas. Each color represents an animal, *N* = 3 mice. Abbreviations: AAA, anterior amygdalar area; Amy, amygdala; AUDp, primary auditory area; BLA, basolateral amygdalar nucleus; BMA, basomedial amygdalar nucleus; CEAl, central amygdala lateral portion; COA, cortical amygdalar area; ECT, ectorhinal cortex; ENTl, entorhinal area; EPd, endopiriform nucleus dorsal part; GPe, globus pallidus external segment; HY, hypothalamus; IA, intercalated amygdalar nucleus; LA, lateral amygdala; NDB, diagonal band nucleus; NLOT, nucleus of the lateral olfactory tract; PER, perirhinal area; PIR, piriform area; RN, raphe nucleus; STRp, posterior striatum; TH, thalamus; cl = contralateral.

To assess whether the large shock-evoked astrocytic Ca²⁺ transients could be explained by movement-related dynamics, we correlated ΔF/F during the shock period with locomotor speed. Maximum shock-evoked Ca²⁺ signals showed only weak correlation with speed (Pearson’s *r* = 0.176; Extended Data Fig. 4c), and cross-correlation analysis revealed a 2.8 s lag of peak correlation between speed and ΔF/F signals (Extended Data Fig. 4d,e), suggesting that movement alone is unlikely to account for the observed Ca^2+^ dynamics.

To directly compare the temporal dynamics of astrocytic and neuronal activity in the TeA, we performed dual-color fiber photometry by co-injecting AAV-hGFAP-GCaMP6f together with AAV-hSyn1-dlox-PinkyCaMP and AAV-mCaMKIIα-iCre into the TeA (Fig. 4e). This strategy allowed simultaneous recordings of astrocytic and excitatory neuronal Ca^2+^ dynamics within the same mice. Both astrocytes and neurons exhibited stimulus-evoked Ca²⁺ signals during fear conditioning and retrieval (Fig. 4f,g). However, peak astrocytic Ca²⁺ transients were significantly delayed relative to neuronal Ca²⁺ signals following CS^+^ and CS^-^ presentation (*p* ≤ 0.01). In addition, neuronal and astrocytic Ca²⁺ dynamics differed significantly at freezing onset (*p* = 0.025; Extended Data Fig. 4f). Thus, neuronal Ca²⁺ dynamics showed significantly more event-locked modulation aligned to freezing behavior than astrocytic Ca²⁺ dynamics. Moreover, average neuronal Ca^2+^ transients during CS^+^ showed high correlation with freezing levels across mice (Pearson’s *r* = 0.857), whereas astrocytic Ca^2+^ transients during CS^+^ showed no correlation with freezing (Pearson’s *r* = 0.096, Extended Data Fig. 4g). These findings indicate that astrocytic Ca²⁺ signaling in the TeA temporally follows neuronal activation and is more closely associated with sensory inputs than with locomotor activity and fear memory. This suggests that global astrocytic Ca^2+^ dynamics are required to support sensory processing of CS^+^ without directly providing one-to-one instructions for fear memory expression.

Whereas the previous results suggest that local neuronal signaling may partially drive astrocytic Ca²⁺ dynamics, long-range projections represent additional potential sources for astrocytic GPCR modulation. The TeA is known to receive inputs from multiple cortical areas (Oh et al., 2014; Zingg et al., 2014), whereas its subcortical afferents remain less well defined. To identify additional subcortical inputs to the TeA, we performed retrograde tracing using cholera toxin subunit B (CTB) injections into the TeA (Fig. 4h). Retrograde labeling identified neurons in multiple subcortical regions, with the strongest contributions arising from the thalamus, cortical amygdalar area, lateral amygdala, basolateral amygdalar nucleus and basomedial amygdalar nucleus (Fig. 4i,j). Notably, several of these regions have been implicated in threat-related processing and value encoding, positioning them as candidate modulators of TeA-dependent fear memory retrieval (Dalmay et al., 2019; Menegas et al., 2018; Soares-Cunha et al., 2020).

Together, these results demonstrate that TeA astrocytes show robust Ca²⁺ increases to sensory and aversive stimuli during fear conditioning and retrieval. We identified local neurons and multiple long-range projections as candidate modulators of astrocytic dynamics during fear retrieval.

### Astrocytic Gi-GPCR signaling suppresses sensory-evoked astrocyte signals while preserving responses in TeA output neurons

To determine how astrocytic Gi-GPCR activation influences sensory processing and circuit activity in TeA, we first monitored Ca²⁺ dynamics in astrocytes expressing both AAV-hGFAP-GCaMP6f and AAV-hGFAP-hM4D_i_-mCherry during fear retrieval (Fig. 5a-b). Mice were presented with CS^+^ stimuli before and after i.p. injection of CNO or saline (Fig. 5c). Prior to CNO administration, CS^+^ presentations reliably evoked Ca²⁺ transients in astrocytes. Strikingly, chemogenetic activation of astrocytic Gi-GPCR signaling led to a significant reduction in sound-evoked astrocytic Ca²⁺ signals compared to saline-treated controls (*p* ≤ 0.01; Fig. 5d-f). These findings indicate that astrocytic Gi-GPCR activation suppresses sensory-evoked astrocytic Ca²⁺ dynamics during retrieval. Additionally, baseline fluorescence levels were significantly reduced in CNO-injected animals compared to saline controls (*p* = 0.03; Fig. 5g,h), suggesting a global reduction of astrocytic activity. This may reflect a change in the state of cortical sensory processing. We hypothesize that although astrocytic Ca²⁺ dynamics do not directly control fear expression, a minimum level of sound-evoked astrocytic activity is necessary for normal sensory processing and may help prevent excessive fear memory retrieval.

**Figure 5:**
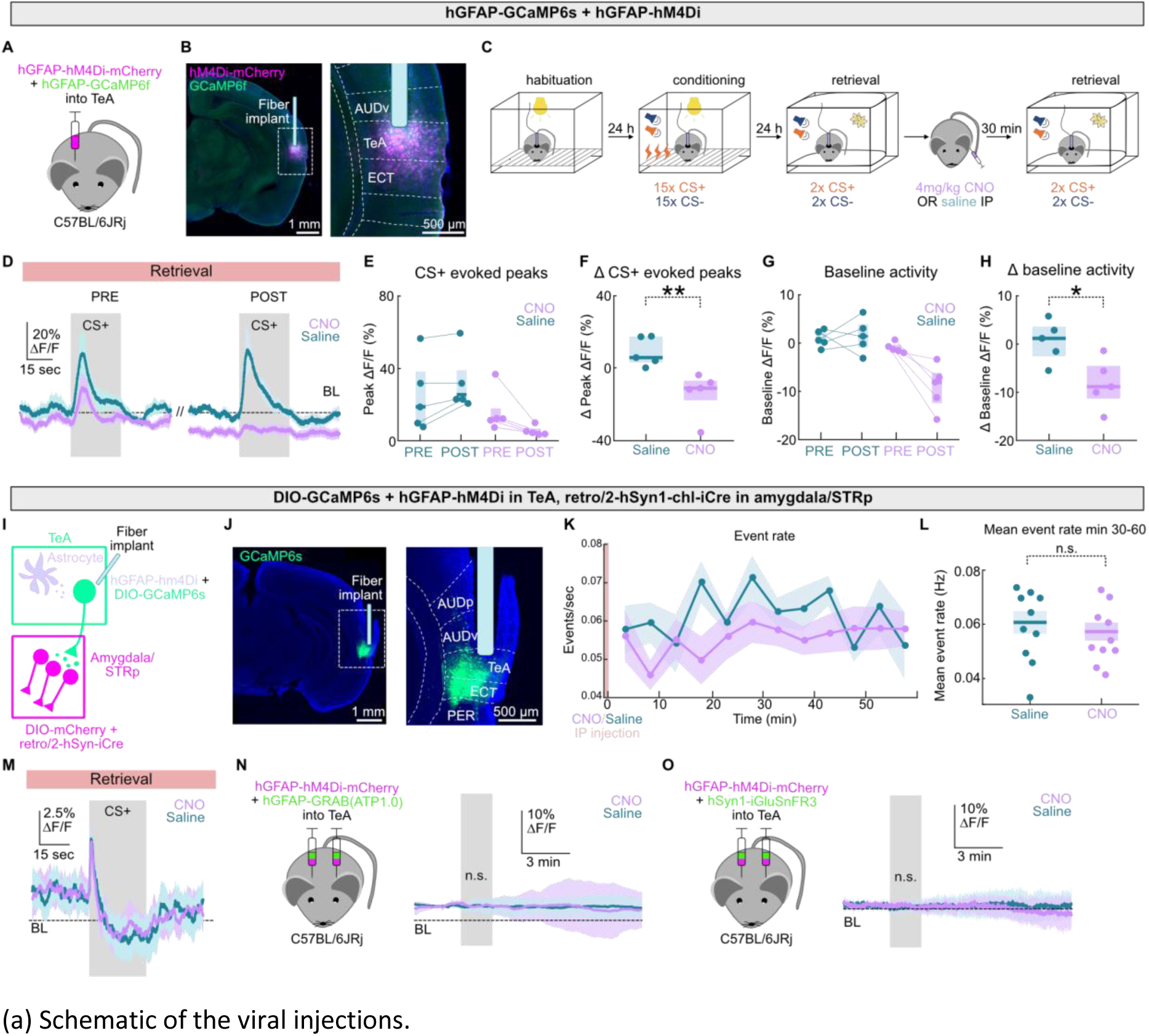
Astrocytic Gi-GPCR signaling suppresses sensory-evoked astrocyte activity while preserving responses in TeA output neurons. (a) Schematic of the viral injections. (b) Exemplary epifluorescence image of injection site and fiber implant. (c) Schematic of behavioral paradigm. (d) Normalized fluorescence changes (ΔF/F) from astrocytes expressing AAV-hGFAP-GCaMP6f and AAV-hGFAP-hM4D_i_-mCherry aligned to CS⁺ presentations before (PRE) and after (POST) i.p. injection of CNO or saline during retrieval, *N* = 5 mice per condition. (e) Peak astrocytic Ca²⁺ transients (baseline-subtracted) upon CS⁺ presentations before (PRE) and after (POST) i.p. injection of saline or CNO. PRE saline: 32.80% [11.86-50.39%] ΔF/F, POST saline: 41.70% [29.37-51.31%] ΔF/F; PRE CNO: 20.93% [13.38-24.22%] ΔF/F, POST CNO: 6.67% [4.40-8.18%] ΔF/F, *N* = 5 mice per condition. (f) Difference in peak astrocytic Ca²⁺ transients upon CS⁺ (POST-PRE) following i.p. injection of saline or CNO; saline: 5.69% [2.95-17.37%]; CNO: –11.30% [-17.96-(–7.15%)], (*p* ≤ 0.01; Mann-Whitney *U* test), *N* = 5 mice per condition. (g) Baseline astrocytic activity before (PRE) and after (POST) i.p. injection of saline or CNO. PRE saline: 0.58% [-0.23-2.48%] ΔF/F, POST saline: 1.43% [-0.92-3.88%] ΔF/F; PRE CNO: –1.32% [-1.68-(–0.36%)] ΔF/F, POST CNO: –8.12% [-12.42-(–6.16%)] ΔF/F, *N* = 5 mice per condition. (h) Difference in baseline astrocytic activity; saline: 1.21% [-2.50-3.62%]; CNO: –8.80% [-11.25-(–4.49%)], (*p* = 0.03; Mann-Whitney *U* test), *N* = 5 mice per condition. (i) Schematic of the experimental strategy used to record Ca²⁺ transients from TeA-LA/STRp-projecting neurons while chemogenetically activating Gi-GPCR signaling in TeA astrocytes. (j) Exemplary epifluorescence images showing AAV-CBA-DIO-GCaMP6s expression in TeA. (k) Event rate of synchronized Ca²⁺ transients in TeA-LA/STRp-projecting neurons over 1 h following i.p. injection of CNO or saline. Injection time is indicated by a peach-colored vertical bar, *N* = 9 mice per condition. (l) Mean synchronized event rate during minutes 30-60 post-injection for CNO- and saline-treated animals (CNO: 0.06 ± 0.01 Hz; saline: 0.06 ± 0.01 Hz, *p* = 0.529, unpaired *t*-test), *N* = 9 mice per condition. (m) Normalized fluorescence changes (ΔF/F) from TeA-LA/STRp-projecting neurons aligned to CS⁺ presentations during retrieval (peak ΔF/F for CNO: 6.51 ± 0.53%; saline: 6.75 ± 0.29%), *N* = 9 mice per condition. (n) Astrocytic transients in acute TeA slices expressing AAV-hGFAP-hM4D_i_-mCherry and AAV-hGFAP-GRAB(ATP1.0). CNO/saline application is indicated by the grey shaded area. No significant difference was detected (peak ΔF/F CNO: 5.04 ± 0.81%, saline: 3.47 ± 0.36%; *p* = 0.481; unpaired *t*-test), *N* = 6 slices per condition. (o) Astrocytic transients in acute TeA slices expressing AAV-hGFAP-hM4D_i_-mCherry and AAV-hSyn1-iGluSnFR3. CNO/saline application is indicated by the grey shaded area. No significant difference was detected when comparing peak responses (peak ΔF/F CNO: 1.05 ± 0.18%, saline: 1.19 ± 0.24%; *p* = 0.854; unpaired *t*-test), *N* = 6 slices per condition. Abbreviations: AUDp, primary auditory area; AUDv, ventral auditory area; ECT, ectorhinal area; PER, perirhinal area; STRp: posterior striatum.

We next asked whether astrocytic Gi-GPCR activation modulates TeA outputs involved in fear memory retrieval. Specifically, we focused on projections to the lateral amygdala (LA) and posterior striatum (STRp), which have established roles in fear-related processing (Cheng et al., 2025; Dalmay et al., 2019). To this end, we labeled TeA neurons projecting to the LA and STRp using AAV2-retro-hSyn1-iCre injections into the target areas combined with Cre-dependent AAV-CBA-DIO-GCaMP6s injections into TeA, while simultaneously expressing AAV-hGFAP-hM4D_i_-mCherry in TeA astrocytes (Fig. 5i,j, Extended Data Fig. 5a). Following CNO administration, Ca²⁺ event rates in TeA-LA/STRp-projecting neurons showed a trend toward lower values compared with controls post-injection (Fig. 5k, Extended Data Fig. 5b). However, no statistically significant difference was detected (Fig. 5l, Extended Data Fig. 5c). Additionally, Ca²⁺ dynamics upon CS^+^ presentations during fear conditioning (Extended Data Fig. 5d), or retrieval (Fig. 5m), freezing-evoked Ca^2+^ dynamics during retrieval (Extended Data Fig. 5e), and mean ΔF/F during retrieval (Extended Data Fig. 5f,g) were not affected. Likewise, TeA-LA/STRp–projecting neurons did not show altered Ca²⁺ dynamics in acute slices upon astrocytic Gi-GPCR activation (Extended Data Fig. 5h). This suggests that other neuronal populations may be the primary targets of astrocytic Gi-GPCR modulation.

Finally, we asked whether astrocytic Gi-GPCR activation induces detectable gliotransmitter release. To monitor extracellular ATP and glutamate levels, we expressed AAV-hGFAP-GRAB(ATP1.0) or AAV-hSyn1-iGluSNFR3 in TeA, either alone or together with AAV-hGFAP-hM4D_i_-mCherry. Bath application of ATP and glutamate evoked robust fluorescence increases in AAV-hGFAP-GRAB(ATP1.0) (Extended Data Fig. 5i) and AAV-hSyn1-iGluSNFR3 expressing slices (Extended Data Fig. 5j), respectively, validating sensor functionality. However, bath-applied CNO did not detectably increase ATP or glutamate signals (Fig. 5n,o). Likewise, bath application of ligands for the major identified endogenous astrocyte GPCRs (Fig. 1; NE, NTS, PACAP, or glutamate) did not evoke measurable ATP or glutamate release (Extended Data Fig. 5k,l).

Together, these findings indicate that while Gi-GPCR activation suppresses sound-evoked astrocytic Ca²⁺ signals, stimulus-locked dynamics in TeA output neurons remain largely intact. This suggests that astrocytes influence fear retrieval through more complex network interactions rather than by directly shaping cue-evoked neuronal signals in output neurons.

### Appetitive auditory learning is unaffected by astrocytic Gi-GPCR activation in TeA

Given that astrocytic Gi-GPCR activation strongly reduced sound-evoked signals, we next asked whether this signaling pathway contributes more generally to auditory learning or is selectively involved in aversive auditory memories.

To address this, we trained mice in an auditory go/no-go task in which one sound predicted food reward (CS^+^) while responses to a second sound (CS^-^) had to be withheld (Fig. 6a). Although frequency-modulated sweeps were used in the aversive conditioning paradigm, pilot experiments indicated that mice poorly discriminated these stimuli in the appetitive task. Therefore, pure tones were used to ensure robust learning.

**Figure 6:**
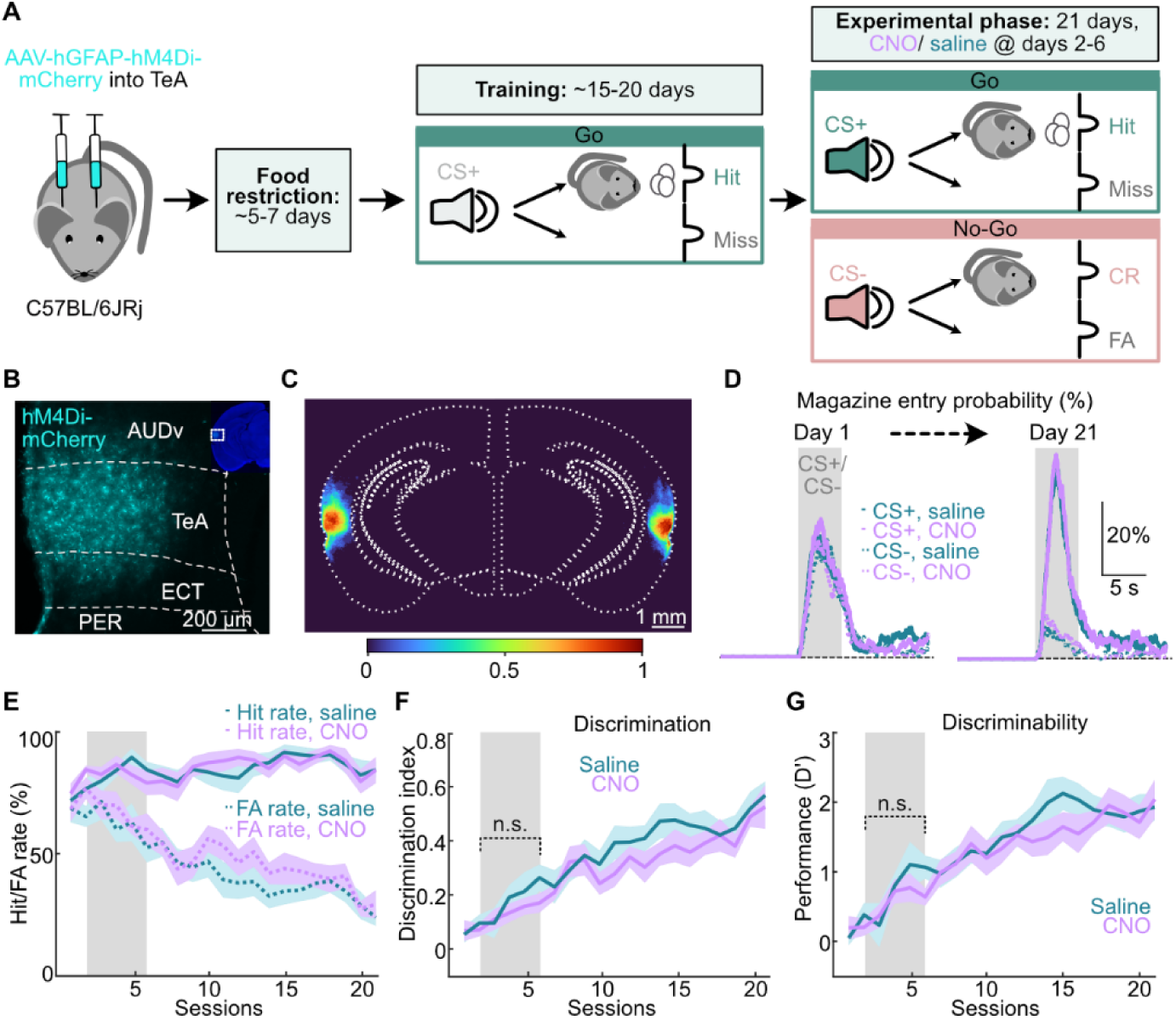
Appetitive auditory learning is unaffected by astrocytic Gi-GPCR activation. (a) Experimental timeline for the auditory go/no-go task. Mice received bilateral AAV-hGFAP-hM4D_i_-mCherry injections into TeA, followed by food restriction, training, and an experimental phase with CNO or saline injections during sessions 2-6. CR, correct rejections; FA, false alarms. (b) Exemplary epifluorescence image of injection site. (c) Overlay of main viral injection sites for AAV-hGFAP-hM4D_i_-mCherry in TeA, for mice used in behavioral tests. Color scale indicates proportion of mice with detectable virus expression. (d) Mean probability of magazine entries aligned to CS^+^ and CS^-^ presentations on session 1 (left) and session 21 (right) of the experimental phase in mice expressing AAV-hGFAP-hM4D_i_-mCherry, treated with CNO or saline. CS^+^ response probability increased and CS^-^ response probability decreased over time, *N* = 11 mice per condition. (e) Hit rate and FA rate across sessions in saline- and CNO-treated mice expressing AAV-hGFAP-hM4D_i_-mCherry, *N* = 11 mice per condition. (f) Discrimination index across sessions for saline- and CNO-treated mice expressing AAV-hGFAP-hM4D_i_-mCherry. (CNO: 0.12 ± 0.03; saline: 0.16 ± 0.04; *p* = 0.695, unpaired *t*-test), *N* = 11 mice per condition. (g) Discriminability (D’) across sessions in saline- and CNO-treated mice expressing AAV-hGFAP-hM4D_i_-mCherry. (CNO: 0.55 ± 0.03; saline: 0.72 ± 0.05; *p* = 0.695, unpaired *t*-test), *N* = 11 mice per condition. Abbreviations: AUDv, ventral auditory area; ECT, ectorhinal area; PER, perirhinal area; TeA, temporal association area.

To investigate the contribution of astrocytic Gi-GPCR signaling to appetitive learning and memory, we expressed AAV-hGFAP-hM4D_i_-mCherry in TeA astrocytes (Fig. 6b,c) and administered CNO or saline during the early learning phase (sessions 2-6). Across the 21 sessions of the experimental phase, both groups showed progressive improvements in task performance, reflected by increased hit rates, reduced false alarm (FA) rates and rising discrimination indices (Fig. 6d-f), indicating successful task acquisition.

Hit rates and FA rates evolved similarly in saline- and CNO-treated mice (Fig. 6e). Consistent with this, astrocytic Gi-GPCR activation did not affect discrimination index (*p* = 0.695; Fig. 6f) or discriminability (*p* = 0.695; Fig. 6g) during the manipulation window (sessions 2-6). Additional behavioral measures, including hit latency and baseline magazine entries, were also indistinguishable between groups (Extended Data Fig. 6a,b). To exclude potential off-target effects of CNO, we repeated the go/no-go paradigm in naïve wild-type mice. CNO treatment did not affect discrimination index or discriminability (both *p* = 0.199; Extended Data Fig. 6c-e), confirming that systemic CNO does not confound behavioral readouts in this paradigm.

Finally, we tested whether TeA glutamatergic neurons contribute to appetitive auditory learning. To this end, hM4D_i_ was expressed in mCaMKIIα^+^ neurons using viral injections of AAV-hSyn1-dlox-hM4D_i_-mCherry together with mCaMKIIα-iCre (Extended Data Fig. 6f). CNO treatment did not alter hit or FA rates (Extended Data Fig. 6g), nor did it affect discrimination and discriminability (both *p* = 1; Extended Data Fig. 6h,i), indicating that neither astrocytic Gi-GPCR signaling nor excitatory neuronal Gi-GPCR signaling in TeA is sufficient to modulate performance during the early acquisition phase of this appetitive auditory task.

Together, these findings indicate that astrocytic Gi-GPCR signaling in TeA modulates aversive memory retrieval, while leaving appetitive auditory learning and memory intact.

## Discussion

Here, we demonstrate that astrocytic Gi-GPCR signaling in the temporal cortex is sufficient to enhance auditory fear memory retrieval. This effect is cell type-specific, GPCR-class-specific, and behaviorally selective, identifying astrocytic Gi signaling as a candidate regulator of how learned sensory cues drive defensive behavior.

Previous studies have identified a critical role for temporal cortex neurons in fear memory retrieval by selectively inhibiting neuronal subpopulations using optogenetic and chemogenetic approaches (Cheng et al., 2025; Dalmay et al., 2019). However, the function of astrocytes in the temporal cortex was still unknown. In some brain areas, activation of astrocytic Gi-coupled receptors enhances synaptic plasticity and improves memory-related behavioral performance (Nam et al., 2019; Robin et al., 2018), whereas another study showed that astrocytic Gi activation disrupts remote, while sparing recent memory (Kol et al., 2020). Thus, astrocytes can bidirectionally regulate memory depending on circuit, timing, and downstream gliotransmitter pathways. Our findings reveal for the first time that astrocytic Gi-GPCR activation in the temporal cortex increases freezing during fear memory retrieval while leaving reward-based learning and memory intact. The absence of an effect during appetitive go/no-go learning indicates that astrocytic Gi-GPCR signaling does not generally enhance auditory association learning. Instead, astrocytic Gi-GPCR signaling appears to preferentially influence cue processing in threat-related contexts, consistent with a model where threat and reward associations lead to distinct restructuring of cortical responses to sensory cues.

Importantly, our data indicate that distinct GPCR classes engage distinct astrocytic signaling programs with divergent functional consequences (Durkee et al., 2019; Oe et al., 2020; Vaidyanathan et al., 2021). While Gq-coupled receptors primarily drive IP_3_-mediated Ca^2+^ release in astrocytes (Araque et al., 2014; Navarrete & Araque, 2010), Gs- and Gi-coupled pathways are associated with cAMP-dependent signaling dynamics (Durkee et al., 2019; Oe et al., 2020; Sun et al., 2013). Importantly, our data and previous studies show that Gi-GPCR signaling in astrocytes can additionally modulate Ca^2+^ release with brain-area-specific downstream consequences (Durkee et al., 2019). In some systems, Gi activation has been linked to increased gliotransmission (Durkee et al., 2019; Y. Li et al., 2020), whereas our data do not support Gi-GPCR-induced gliotransmission. Instead, our findings are consistent with the idea that gliotransmission may require strong and sustained Ca^2+^ signaling (H. Li et al., 2025), which may not be achieved under our experimental conditions. Thus, our data highlight that molecular divergence of astrocytic GPCR signaling translates into functional divergence at the behavioral level. Such functional heterogeneity in astrocytic GPCR signaling has been observed previously in the regulation of sleep, where Gi- and Gq-coupled signaling exert distinct, and even opposing, effects on brain state (Vaidyanathan et al., 2021).

Moreover, our data suggest that temporal cortical astrocytes are primarily driven by sensory input rather than by fear state or mnemonic functions. Thus, astrocytes respond robustly to both auditory cues and foot shocks, yet cue-evoked Ca^2+^ responses do not increase across learning and do not correlate with freezing during retrieval. This contrasts with recent findings in the basolateral amygdala, where astrocytes dynamically track fear-related states and support neural representations of fear memory (Bukalo et al., 2026; Ghenissa et al., 2026; Suthard et al., 2023). At the same time, strong sensory-evoked astrocyte responses have been reported in primary sensory cortices (Paukert et al., 2014; Wang et al., 2006). These findings point to regional specialization of astrocyte function, with cortical astrocytes predominantly reflecting sensory input, whereas subcortical astrocytes may more directly encode behavioral or emotional state.

Of note, we show that astrocytic hM4D_i_ activation reduces Ca^2+^ responses during sensory cue presentation in the temporal cortex. The slow time scale of hM4D_i_-induced processes likely excludes rapid event-specific effects. Instead, Gi signaling is expected to shift the gate or gain for responses to faster signals. Given the non-linear nature of astrocytic Ca^2+^ release, even a sustained hM4D_i_-induced signal can be expected to disproportionately reduce prominent sensory-evoked responses. This aligns with previous work showing that neuromodulatory GPCR signaling can gate astrocyte responsiveness rather than merely scaling baseline activity (Guttenplan et al., 2025; Paukert et al., 2014). In these studies, noradrenergic signaling enables astrocytic responses to other neurotransmitters and sensory input. Similarly, the neuromodulator histamine increases the likelihood of global Ca^2+^ events (Taylor et al., 2025). Here, we show for the first time that astrocytic Gi-GPCR signaling exerts gating-like effects on sensory representations during retrieval. Surprisingly, decreased astrocytic Ca^2+^ transients were paralleled by improved fear memory retrieval, suggesting that astrocytic Ca^2+^ transients naturally suppress fear responses. Astrocytic Gi-GPCR signaling may thus reshape how sensory cues are translated into defensive behavior.

Future studies should address several limitations of this study. First, it remains unclear to what extent chemogenetic activation replicates the subcellular spatiotemporal dynamics of endogenous GPCR signaling. Thus, directly modulating endogenous GPCRs will be key for future studies to determine whether endogenous Gi-GPCR activation similarly increases fear memory retrieval. Second, recent transcriptomic work has identified distinct astrocyte subtypes in the auditory cortex, suggesting potential differences in GPCR expression and signaling across subpopulations (Chai et al., 2017; Gungor Aydin et al., 2024). Direct manipulation of neuromodulator-specific GPCRs will be required to resolve this question. Third, fiber photometry lacks subcellular resolution and may underestimate localized astrocytic signaling events and activity restructuring in neuronal populations. Follow-up studies using two-photon imaging will be necessary to reveal to what extent localized Ca^2+^ signaling and gliotransmitter release contribute to the observed effects. Fourth, the downstream neuronal targets and circuit mechanisms through which TeA astrocytes influence retrieval remain unknown. Future work should determine whether altered astrocytic processing of sensory cues changes potassium buffering, modulation of cortical blood flow, or other forms of astrocyte-to-neuron signaling in behaviorally relevant projection neurons.

Together, our findings identify astrocytic Gi-GPCR signaling in the temporal cortex as a powerful regulator of auditory fear memory retrieval. Notably, endogenous astrocytic Ca^2+^ transients appear to primarily reflect sensory cue processing independent of behavioral output. We propose a model where a basal tone of astrocytic Ca^2+^ signaling is required to exert a suppressive modulation of fear memory expression without providing precise one-to-one instructions for the amount of memory retrieval. Thus, our data point to more complex astrocyte-neuron interactions than previously thought. Investigating how localized, synapse-specific Ca^2+^ dynamics, astrocytic mechanisms beyond Ca^2+^ signaling, and additional astrocyte-neuron interactions contribute to fear memory retrieval will provide additional insights into underlying mechanisms. Together, our results provide a framework for understanding how neuromodulator-responsive astrocytes influence emotional memory retrieval through receptor-specific signaling mechanisms.

## Methods

### Animals

Unless otherwise stated, C57BL/6JRj (Janvier; RRID:IMSR_RJ:C57BL-6JRJ) and the following Cre-driver lines were used in this study: *Sst-IRES-Cre* (B6J.Cg-Ssttm2.1(cre)Zjh/MwarJ; The Jackson Laboratory, RRID:IMSR_JAX:028864), *Vip-IRES*-Cre (B6J.Cg-VIPtm1(cre)Zjh/AreckJ; The Jackson Laboratory, RRID:IMSR_JAX:031628), and *Pvalb-IRES-Cre* (B6.129P2-Pvalbtm1(cre)Arbr/J; The Jackson Laboratory, RRID:IMSR_JAX:017320). All transgenic lines were backcrossed to C57BL/6JRj mice for at least 7 generations.

Mice were housed in enriched standard laboratory cages under a 12 h light/dark cycle, with *ad libitum* access to food and water, unless otherwise stated. Animals used for *in vitro* experiments were group-housed, whereas those assigned to behavioral testing were single-housed beginning on the second day of handling. Female mice were used for behavioral experiments, and both sexes were used for all other experiments.

All procedures complied with the institutional guidelines of the Medical University of Vienna and were approved by the Austrian Ministry of Science.

### Surgical procedures

Stereotaxic surgeries were performed on mice aged 6-12 weeks. Anesthesia was induced using 5% isoflurane and maintained at 1-2.5% throughout the procedure. Body temperature was kept at 37 °C using a feedback-controlled heating pad. Prior to incision, lidocaine (3.5 mg/kg^-1^; Gebro Pharma, cat. no. 100562) and carprofen (4 mg/kg^-1^; Zoetis, cat. no. 256684) were administered subcutaneously for local anesthesia and general analgesia. Bregma and lambda were aligned in a stereotactic frame (David Kopf Instruments).

For targeting the temporal association area (TeA), a small craniotomy was made to perform injections at –3.6 mm posterior (AP) relative to bregma, 0.35 mm medial to the lateral skull ridge (ML) and –1.23 mm dorsoventral (DV) from the pial surface. For targeting the amygdala/posterior striatum, injections were made at –0.3 mm AP, 3.9 mm ML from the midline, and –3.9 mm DV using a 25° posterior angle and 20° lateral rotation of the injector. In behavioral experiments involving chemogenetic manipulations and in slice imaging experiments, injections were administered bilaterally in the TeA; in experiments involving fiber photometry, they were administered unilaterally.

Viral vectors were delivered using a glass micropipette at 40 nL/min, with injected volumes ranging from 30-200 nL. For chemogenetic manipulation of astrocytes in behavioral experiments, C57BL/6JRj mice received 100 nL of ssAAV-5/2-hGFAP-hM4D(Gi)_mCherry-WPRE-hGHp(A) (2.2*10^12 gc/mL; VVF UZH, cat. no. v103-5), 50-100 nL of ssAAV-5/2-hGFAP-hM3D(Gq)_mCherry-WPRE-hGHp(A) (0.5-1.2*10^12 gc/mL; VVF UZH, cat. no. v97-5) or 75 nL of ssAAV-5/2-hGFAP-HA_rM3D(Gs)_IRES_mCitrine-WPRE-hGHp(A) (3.2*10^12 gc/mL; VVF UZH, cat. no. v97-5).

For neuronal chemogenetic manipulation, 70-100 nL of ssAAV-9/2-hSyn1-dlox-hM4D(Gi)_mCherry(rev)-dlox-WPRE-hGHp(A) (7.7-12*10^11 gc/mL; VVF UZH, cat. no. v84-9 (Krashes et al., 2011)) was injected into *Sst-IRES-Cre*, *Vip-IRES-Cre*, or *Pvalb-IRES-Cre* mice, or co-injected with ssAAV-9/2-mCaMKIIα-iCre-WPRE-hGHp(A) (6.6*10^10 gc/mL; VVF UZH, cat. no. v206-9) into C57BL/6JRj mice. For fiber photometry experiments, 200 nL of ssAAV-5/2-hGFAP-hHBbl/E-GCaMP6f-bGHp(A) (5.6*10^12 gc/mL; VVF UZH), AAV-9/2-CBA-DIO-GCaMP6s-P2A-mBeRFP-WPRE-pA (2.9*10^12 gc/mL; VVF UZH, cat. no. v828-9 (Melzer et al., 2021)), or ssAAV-9/2-hSyn1-dlox-NES_6xHis_PinkyCaMP(rev)-dlox-WPRE-bGHp(A) (6.8*10^12 gc/mL; VVF UZH, cat. no. v1125-9 (Fink et al., 2024)) were injected into the TeA of C57BL/6JRj mice. For projection-specific photometry or slice imaging experiments, 200 nL of ssAAV2-retro-hSyn1-chl-iCre-WPRE-SV40p(A) (7.9*10^12 gc/mL; VVF UZH, cat. no. v223-retro) was co-injected with ssAAV-9/2-hSyn1-DIO-mCherry(rev)-DIO-WPRE-hGHp(A) (1*10^12 gc/mL; VVF UZH, cat. no. v116-9) into the amygdala/posterior striatum. For other slice imaging experiments, 200 nL of ssAAV-5/2-hGFAP-hHBbl/E-GCaMP6f-bGHp(A) (5.6*10^12 gc/mL; VVF UZH, cat. no. v275-5), 100 nL of ssAAV-DJ/2-hSyn1-chI-iGluSnFR3.v857.GPI(c.-o.)-WPRE-bGHp(A) (1*10^12 gc/mL; VVF UZH, cat. no. v1039-DJ), or 200 nL ssAAV-5/2-hGFAP-chI-GRAB(ATP1.0)-WPRE-hGHp(A) (7*10^12 gc/mL; VVF UZH, cat. no. v1170-5 (Wu et al., 2022)) was injected into the TeA.

The injection pipette was left at the injection site for 3 min before and after infusion to minimize reflux. For fiber photometry experiments, the skull skin was removed, and the skull surface was roughened to improve adhesion. Mono fiberoptic cannulae (Doric Lenses, cat. no. MFC_200/230–0.37_3mm_MF1.25_A45) were implanted at the injection site at –1.23 DV, with the angled fiber tip facing anterior. The craniotomy was sealed with Kwik-cast (World Precision Instruments), and the implant was secured with cyanoacrylate glue reinforced with Paladur (Kulzer, cat. no. L010131) and covered with black nail polish to reduce ambient light scatter.

Animals recovered in a heated recovery cage and were monitored for 4 consecutive days. Carprofen (4 mg/kg) was administered for 2 days post-surgery. Experiments began 3-4 weeks after surgery to allow for viral expression.

### Plasmids

pAAV-GFAP-hM4D(Gi)-mCherry was a gift from Bryan Roth (Addgene plasmid # 50479; http://n2t.net/addgene:50479; RRID:Addgene_50479).

pAAV-GFAP-hM3D(Gq)-mCherry was a gift from Bryan Roth (Addgene plasmid # 50478; http://n2t.net/addgene:50478; RRID:Addgene_50478).

pAAV-GFAP-HA-rM3D(Gs)-IRES-mCitrine was a gift from Bryan Roth (Addgene plasmid # 50472; http://n2t.net/addgene:50472; RRID:Addgene_50472).

pAAV-hSyn-DIO-hM4D(Gi)-mCherry was a gift from Bryan Roth (Addgene plasmid # 44362; http://n2t.net/addgene:44362; RRID:Addgene_44362).

pAAV-mCaMKIIα-iCre-WPRE-hGHp(A) was constructed by the VVF (iCre: Addgene #24593).

pAAV-hGFAP-hHBbl/E-GCaMP6f-bGHp(A) was constructed by the VVF (GCaMP6f: Addgene #51083; hGFAP promoter fragment: DOI: 10.1002/glia.20622).

pAAV-CBA-DIO-GCaMP6s-P2A-mBeRFP-WPRE-pA was a gift from Bernardo Sabatini (Addgene plasmid # 175177; http://n2t.net/addgene:175177; RRID:Addgene_175177).

pssAAV-2-hSyn1-dlox-NES_6xHis_PinkyCaMP(rev)-dlox-WPRE-bGHp(A) was provided by Olivia Masseck, University of Cologne, Germany, and corresponds to Addgene #232857.

pAAV-hSyn1-chl-iCre-WPRE-SV40p(A) was constructed by the VVF (iCre: Addgene #24593).

pAAV-hSyn-DIO-mCherry was a gift from Bryan Roth (Addgene plasmid # 50459; http://n2t.net/addgene:50459; RRID:Addgene_50459).

pAAV-hSyn1-chI-iGluSnFR3.v857.GPI(c.-o.)-WPRE-bGHp(A) was constructed by the VVF (iGluSnFR3.v857.GPI(c.-o.): Addgene #175181).

pAAV-GfaABC1D-GRAB_ATP1.0 was a gift from Yulong Li (Addgene plasmid # 167579; http://n2t.net/addgene:167579; RRID:Addgene_167579).

### Perfusion and tissue preparation

At the end of behavioral and histological experiments, mice were deeply anesthetized with 5% isoflurane and transcardially perfused with 4% paraformaldehyde (PFA; Biotrend Chemikalien, cat. no. 30400000-2) in phosphate-buffered saline (PBS). Brains were extracted and sectioned coronally at 50 µm for immunohistochemistry and 100 µm for all other analyses using a vibratome (Leica VT1000S) 24h post-fixation. Sections were either mounted directly using medium containing DAPI (Fisher Scientific, cat. no. 15395816) onto adhesion slides or further processed for immunohistochemistry. Images were acquired using a Vectra Polaris imaging system (Akoya Biosciences).

### Histological overviews

Viral expression was plotted based on mCherry or mCitrine-expression in coronal sections using a custom-written MATLAB script (MathWorks, version R2022a). The sections were aligned, background-subtracted, binarized and overlaid across animals for the main injection site. For each pixel, expression probability was calculated as the fraction of animals showing signal at that location, generating heatmaps scaled from 0 to 1.

### Immunohistochemistry

For immunohistochemistry, free-floating brain slices were rinsed three times for 5 min in PBS and incubated for 1 h at room temperature in a blocking solution containing 5% normal goat serum (NGS, Abcam, cat. no. ab7481) and 0.1% Triton X-100 (Sigma-Aldrich, cat. no. T92384) in PBS. Sections were then incubated with primary antibodies diluted in fresh blocking solution for 24 h at 4°C on a shaking platform.

Following primary antibody incubation, sections were washed five times for 5 min in PBS containing 0.1% Triton X-100, followed by incubation with species-appropriate secondary antibodies diluted in blocking solution for 1 h at room temperature. Finally, the slices were rinsed twice for 5 min in PBS containing 0.1% Triton X-100 and twice in PBS before mounting onto adhesion slides.

The following primary antibodies were used: rabbit anti-GFAP (1:1000; Agilent, cat. no. Z03349-2) and chicken anti-mCherry (1:2000; Fisher Scientific, cat. no. NBP2-25158). Secondary antibodies included goat anti-rabbit IgG (H+L) conjugated to Alexa Fluor 647 (1:500; Fisher Scientific, cat. no. 10729174), goat anti-chicken IgG (H+L) conjugated to Alexa Fluor 488 (1:500; Fisher Scientific, cat. no. A11039), and goat anti-rabbit IgG (H+L) conjugated to Alexa Fluor 488 (1:500; Fisher Scientific, cat. no. 10123672).

To quantify the astrocytic specificity of hGFAP-driven hM4D_i_ expression, immunohistochemical sections were analyzed for overlap between virally expressed mCherry and endogenous GFAP immunoreactivity. For each animal, one to three sections spanning the main injection site were imaged on a laser scanning confocal microscope (Leica TCS SP8) using a 63 x/1.3 glycerin immersion objective (z-stack with pixel size of 0.48 µm). For mCherry-expressing cells, inclusion required a well-defined fluorescent structure with visible soma or characteristic astrocytic processes spanning multiple focal planes. GFAP-positive astrocytes were identified based on a morphology consistent with astrocytic processes. All mCherry-positive cells and all GFAP-positive cells within the injection area were manually counted, and the number of cells showing overlapping mCherry and GFAP signal was determined. Astrocytic specificity was calculated as the percentage of mCherry-expressing cells that were also GFAP-positive. Values were first averaged within each slice and then across slices to obtain one value per animal.

### Retrograde tracing

For retrograde tracing, 15-40 nL of 1 µg/µL cholera toxin subunit B Alexa Fluor 647 Conjugate (CTB; Fisher Scientific, cat. no. 11574267) or cholera toxin subunit B Alexa Fluor 555 Conjugate (CTB; Fisher Scientific, cat. no. 11584247) was injected into the TeA. Three weeks after the surgery, mice were transcardially perfused, the whole brain was embedded in 4% agarose and coronally sectioned into 50 µm slices as described previously. 15 subcortical regions exhibiting the highest fluorescence density were selected for analysis. To ensure a reliable and representative sampling of the retrograde input, every second coronal section was analyzed across the entire rostro-caudal extension of the brain. Regions of interest (ROIs) were manually delineated in ImageJ by aligning sample slices with the corresponding reference templates from the Allen Brain Institute atlas.

Manual cell quantification was performed using the ImageJ Cell Counter plugin. Retrogradely labeled neuronal somas were identified based on morphological criteria: intensely fluorescent, round, and sharply defined structures, mostly featuring a weakly fluorescent nucleus.

The total number of labeled cells within each brain area was normalized to the total number of subcortical CTB-positive cells summed across all 15 analyzed regions per mouse. Thalamus (TH) includes RE (nucleus reuniens), PVT (paraventricular thalamic nucleus), RH (rhomboid nucleus), SMT (submedial thalamic nucleus), VM (ventromedial nucleus), PR (prethalamic region), MD (mediodorsal thalamus), IMD (intermediodorsal nucleus), VPL (ventral posterolateral nucleus), VPM (ventral posteromedial nucleus), VAL (ventral anterior-lateral complex), RT (reticular thalamic nucleus), PP (peripeduncular nucleus), SPFp+m+d (subparafascicular nucleus; parvicellular + magnocellular + dorsal parts), AV (anteroventral nucleus), AMd+v (anteromedial nucleus; dorsal + ventral parts), CM (central medial nucleus), MGd+v (medial geniculate nucleus; dorsal + ventral), LP (lateral posterior nucleus), PF (parafascicular nucleus).

### RNA sequencing analysis

Publicly available single-cell RNA sequencing datasets from Tasic et al. (2016), Sugino et al. (2019), and Yao et al. (2021) were analyzed with custom-written MATLAB scripts to assess GPCR expression in astrocytes. Data were downloaded from GEO accession numbers GSE71585, GSE185862 and GSE79238. We used original cell cluster assignments as provided in the metadata. For the Tasic et al. and Yao et al. datasets, cells annotated as astrocytes by the original authors were extracted and averaged.

For the Sugino et al. dataset, the Isocortex-Gfap cluster was extracted and averaged. Astrocyte identity was verified by enrichment of *Aqp4, Sox9, Aldh1l1, Gfap,* and low expression of the neuronal marker *Rbfox3*, ependymal marker *Foxj1*, perivascular fibroblast marker *Pdgfra*, glutamatergic markers *Slc17a6* and *Slc17a7,* and GABAergic markers *Gad1* and *Gad2*.

We used spatial transcriptomics data from 4 sagittal mouse slices, including areas such as cortex, hippocampus, striatum, thalamus, olfactory bulb, cerebellum, and brainstem (Xenium, 1 female, 3 male mice). Data from male and female mice were combined because no significant sex differences were detected. Xenium graph-based clustering (GEX) algorithm (Generated by the 10x Genomics Xenium Explorer 4.1.1) detected one to three astrocyte-rich clusters per slice that had high expression of *Aqp4*, *Sox9* and *Aldh1l1*, and low expression of the neuronal marker *Rbfox3*, the ependymal marker *Foxj1* and the perivascular fibroblast marker *Pdgfra*.

All scRNA-seq data were analyzed as log(TPM+1). To improve visualization across different datasets, each dataset was normalized to the average expression across all shown GPCRs (main and supplementary figures). Only genes with data available for at least half of the datasets were included. The heatmap was scaled to the maximum value across all data.

### Spatial transcriptomics

This workflow was adapted from a published protocol (Ma et al., 2024). Adult C57BL/6J (Strain #:000664, RRID:IMSR_JAX:000664) mice of both sexes (15-18 weeks old) were deeply anesthetized with 4% isoflurane and transcardially perfused at approximately 5 mL/min with ice-cold RNase-free 1× PBS for 5 min, followed by ice-cold RNase-free 4% PFA for 5 min. Brains were rapidly dissected in an RNase-free workspace and post-fixed in RNase-free 4% PFA at 4 °C overnight (up to 24 h).

Following fixation, brains were cryoprotected by immersion in 30% sucrose until the tissue sank (minimum 1-2 days). Tissue was stored in 30% sucrose at 4 °C until embedding. For sagittal sectioning, brains were bisected along the midline into hemispheres using a fresh blade in an RNase-free environment. Hemispheres were embedded in OCT in 15 × 22 × 22 mm molds, oriented with the midline facing down, and frozen in a dry ice/ethanol slurry. Embedded blocks were stored at –80 °C until sectioning.

For cryosectioning, tissue blocks were equilibrated in the cryostat (CM 1950, Leica) for at least 1 h prior to cutting. The cryostat chamber and specimen holder were maintained at –20 °C. Sections were cut sagittally at 15 µm thickness. Regional landmarks were verified using the Allen Brain Atlas (Allen Institute for Brain Science, 2011) and Paxinos (Paxinos & Franklin, 2019) to confirm anatomical level and orientation. Using the Allen Brain Atlas as a reference, we selected tissue corresponding to sagittal sections 9 and 17/18, and collected these regions as 15 µm cryosections for Xenium analysis (Janesick et al., 2023).

Cryosections were collected into RNase-free 1× PBS and mounted onto 10x Genomics Xenium-issued slides supplied with the Xenium assay reagents, consumables, and gene panel kit. Tissue sections were floated in RNase-free PBS and transferred onto slides using rounded glass pipettes to minimize tissue tearing. Excess PBS was carefully removed, and slides were allowed to fully air-dry before being placed in slide mailers. Mounted slides were stored at –80 °C until submission for Xenium processing.

After all tissue processing and slide preparation were completed in the laboratory, the slides were submitted to the Human Immune Monitoring Shared Resource (HIMSR) at the University of Colorado Anschutz Medical Campus, a core facility that provides spatial biology imaging services. Xenium 5k assay processing and imaging were performed by HIMSR personnel using the 10x Genomics Xenium platform, in accordance with manufacturer protocols and the facility’s standard operating procedures.

Data from 4 sagittal mouse brain sections were then analyzed using 10x Genomics Xenium platform, Xenium Explorer v4.1.1. Xenium graph-based clustering (GEX) algorithms were used. 1 to 3 astrocyte-rich clusters per slice that had high expression of *Aqp4*, *Sox9* and *Aldh1l1*, and low expression of the neuronal marker *Rbfox3*, the ependymal marker *Foxj1* and the perivascular fibroblast marker *Pdgfra* were extracted. Cells from the identified clusters were distributed across all regions of the sagittal slices. Average transcript counts per identified cluster were extracted for further analysis in MATLAB. For better visualization, all data were normalized to the mean expression level across all analyzed GPCRs.

Across datasets, expression of GPCR genes was compared qualitatively by ranking relative expression levels within astrocyte clusters.

### Epifluorescence slice imaging

Mice were deeply anesthetized with 5% isoflurane and transcardially perfused with 30 mL cooled sucrose-based slicing solution (212 mM sucrose, 3 mM KCl, 1.25 mM NaH_2_PO_4_, 26 mM NaHCO_3_, 1 mM MgCl_2_, 0.2 mM CaCl_2_, 10 mM glucose) saturated with carbogen (95% O_2_ / 5% CO_2_). The brains were rapidly extracted and sectioned into 300-µm-thick coronal slices in cold, carbogenated sucrose solution using a vibratome (Leica VT1200). Slices were transferred to carbogenated Ringer’s solution (125 mM NaCl, 25 mM NaHCO_3_, 1.25 mM NaH_2_PO_4_, 2.5 mM KCl, 2 mM CaCl_2_, 1 mM MgCl_2_, 25 mM glucose) for 10 min at 37 °C and subsequently maintained at room temperature until use.

Recordings were performed on an Olympus BX51WI microscope equipped with an ORCA-Fusion Digital CMOS camera (Hamamatsu, cat. no. C14440-20UP). A 10 x water-immersion objective (Olympus, UMPLFLN10XW) was used for slice imaging and a 40 x objective (Olympus, LUMPLFLN40XW) for electrophysiological recordings. During recordings, slices were continuously perfused with carbogenated Ringer’s solution at ∼32 °C.

For fluorescence imaging, genetically encoded fluorescent sensors (see above) were used. Green fluorescent protein (GFP)-based reporters were recorded using a GFP filter set (AFH, cat. no. F46-002; excitation 470/40 nm, emission 525/50 nm, 495 nm beam splitter T495 LPXR; AFH, cat. no. F48-495). Red fluorescent proteins (RFP) were imaged using an RFP filter set (excitation 575/32 nm, emission 625/32 nm, 604 nm beam splitter; AHF). The first 50 frames of each recording were acquired in the RFP channel to document expression where applicable. The camera shutter and excitation LED were synchronized via Micro-Manager software (D. Edelstein et al., 2014). Images were acquired at 4 Hz with 50-100 ms exposure times.

Regions of clear cellular fluorescence within the TeA were selected for imaging, and a 5-20 min baseline period was recorded to allow equilibration of the slices and reduce steep bleaching curves. Ringer’s solution contained tetrodotoxin (TTX; 500 nM; Tocris, cat. no. 1069/1) to block action potential-dependent neuronal activity.

Experimental recordings lasted 27-28 min. GPCR ligands were bath-applied between 3 and 5 min of the recording of the green fluorophore. The following ligands were used: clozapine N-oxide dihydrochloride (CNO) (10 µM; Tocris, cat. no. 6329/50), norepinephrine (10 µM; targetmol, cat. no. TGM-T0871-50mg), PACAP1-38 (300 nM; Phoenix Pharmaceuticals, cat. no. 052-05), neurotensin (300 nM; Phoenix Pharmaceuticals, cat. no. 048-09), adenosine 5’-triphosphate disodium salt hydrate (ATP; 100 µM; Sigma, cat. no. A7699-1G), L-glutamic acid monosodium salt hydrate (glutamate; 1 mM; Sigma, cat. no. G5889-100G), and endothelin-1 (100 nM; Phoenix Pharmaceuticals, cat. no. 023-01). Control experiments were conducted by infusing matched volumes of double-distilled H_2_O (ddH_2_O). From min 15 to the end of the recording, 30 mM KCl (Fisher Scientific, cat. no. 10735874) was applied to confirm slice viability.

#### Slice imaging data analysis

Data analysis was performed using custom-written MATLAB scripts (MathWorks, version R2022a). Fluorescence signals were extracted from manually defined regions of interest (ROIs) corresponding to individual fluorescent cells or astrocyte “domains” (including cell bodies and processes) within the TeA. Fluorescence changes were expressed as ΔF/F by normalizing fluorescence changes to the mean baseline fluorescence during the first 3 min of recording. To correct for slow fluorescence bleaching during imaging, an exponential decay function was fitted to the lower 25% percentile of the control traces and subtracted from all recordings, thereby removing gradual baseline drift while preserving stimulus-evoked responses. Slices that failed to respond to 30 mM KCl application were excluded from further analysis. For quantification of ligand-evoked responses, fluorescence traces were averaged across all ROIs within a slice to obtain a single representative trace per slice. To reduce the influence of spontaneous fluctuations on peak responses, each slice trace was smoothed using a moving-average filter over a 1-min window. Response amplitude was then defined as the maximum smoothed ΔF/F value occurring within 4-10 min of the recording. Responses were compared against control recordings in which matched volumes of ddH_2_O or saline were bath-applied. For statistical comparisons, ≥ 6 slices from ≥ 3 animals per condition were collected. Data are reported as mean ± SEM across slices unless stated otherwise.

### Auditory fear conditioning

Female mice (*N* = 18-20 per experiment; 8-12 weeks old) underwent stereotaxic AAV injections as described above. To control for behavioral effects of CNO in the absence of Designer Receptors Exclusively Activated by Designer Drugs (DREADD) expression, two independent cohorts of naïve female C57BL/6JRj mice (total *N* = 40, including WT siblings of backcrossed Cre lines) underwent the full auditory fear conditioning paradigm without viral DREADD expression.

After a recovery period of 14-20 days, mice were handled for 7 consecutive days and habituated to the experimenter. From the second day of handling onward, mice were single-housed.

Behavioral experiments were conducted between Zeitgeber time (ZT) 1 and 5 in custom-built conditioning boxes consisting of a shock-grid floor enclosed by acrylic plates (31 x 23.5 x 37 cm; length x width x height). Each box was equipped with a Blackfly S USB3 camera (BFS-U3-13Y3M-C, Teledyne), visible and infrared light sources, and a speaker. Auditory stimuli were played from a Titanium Dome Tweeter (TW025A20 1, Audax) driven by a SparkFun MP3 player shield (12660. SparkFun Electronics) and calibrated to 55 dBA at mouse height. Stimulus presentation, lights and shock delivery were controlled via an Arduino microcontroller and logged using a custom MATLAB script through a T7 DAQ Board (LabJack). Videos were recorded at 30 frames per second.

The fear conditioning paradigm was adapted from Letzkus et al. (2011) and spanned three consecutive days. All sessions were conducted during the early light phase. On day 1 (habituation), mice explored the chamber for 10 min without stimuli. On day 2 (conditioning), a 3 min baseline was recorded, followed by presentation of two complex sounds (upsweeps or downsweeps; 30 repetitions of 500 ms sweeps ranging from 5-10 kHz with 45-180 s interstimulus intervals). 15 repetitions of each sound were delivered in pseudorandom order. One sound (CS^+^) co-terminated with a 1 s foot shock (0.6 mA), whereas the other (CS^-^) remained unpaired. Assignment of upsweeps versus downsweeps as CS^+^ or CS^-^ was counterbalanced across animals. On day 3 (retrieval), mice were weighed and injected intraperitoneally (i.p.) with CNO (4 mg/kg in 0.9% NaCl solution (saline)) or saline (control) 30 min prior to testing for chemogenetic experiments. Contextual cues were altered to reduce contextual fear: the grid floor was covered with white plastic, the chamber was reshaped into a semicircle, the visible light was turned off, and a vanilla odor was introduced. After a 3 min baseline, mice received 15 CS^+^ and 15 CS^-^ presentations in the same order as during conditioning, without shock delivery.

#### Analysis of freezing behavior

Freezing on the retrieval day was quantified using a custom MATLAB script. Freezing was defined as the absence of movement except for respiration for ≥ 2 s. A binary mask of the mouse was generated, and frame-to-frame pixel changes were quantified. Thresholds distinguishing freezing from movement were semi-automatically determined for each video based on presentation of predicted freezing segments with an increasing or decreasing number of pixel changes. Ambiguous freezing segments close to defined thresholds were semi-automatically validated. Freezing in response to CS^+^ and CS^-^ was calculated by averaging freezing levels across all sound presentations (15 trials per stimulus). Freezing values reflect absolute freezing levels and were not baseline-subtracted. Baseline freezing was quantified during the 3 min period preceding stimulus presentation. A discrimination index was calculated as:

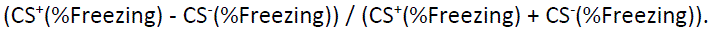

#### Fiber photometry during fear conditioning and retrieval

Mice implanted with fiber-optic cannulae experienced the same behavioral paradigm as described above. During the final two days of handling, mice were habituated to gentle manipulation of the implant and patch cord connection.

Immediately before behavioral testing, implanted fiber-optic cannulae were connected to a 200 µm diameter, 5.5 m long patch cord (NA 0.37; Doric Lenses, MFP_200/220/900–0.37_5.5m_FCM-MF1.25, low autofluorescence epoxy). Excitation light was provided by two fiber-coupled LEDs: a blue LED (470 nm; fiber-coupled LED; Thorlabs; cat. no. M470F3) and a green LED (565 nm; fiber-coupled LED, 9.9 mW; Thorlabs; cat. no. M565F3). The excitation light paths were guided through a Fluorescence MiniCube (Doric lenses; iFMC5-G2_E1(460-490)_F1(500-540)_E2(555-570)_F2(580-680)_S).

Excitation light power at the patch cord tip was set to 45 µW. Signals were detected and amplified using Doric’s built-in photodetectors and amplifiers, acquired using a T7 Daq Board (LabJack), and recorded with custom MATLAB software. LED driver output was sinusoidally modulated at 171 Hz (blue excitation) and 228 Hz (green excitation), and data were sampled at 2052 Hz as described previously by Melzer et al. (2021) and Owen & Kreitzer (2019).

Fluorescence signals were demodulated by extracting the spectral power at the respective modulation frequencies from a short-time Fourier transform computed in MATLAB using a 200-sample sliding window and 180-sample overlap. To correct for background autofluorescence originating from the optical setup and patch cord, fluorescence power was measured prior to connecting the patch cord to the implanted fiber and subtracted from all recorded signals before further analysis. Photobleaching was corrected by fitting an exponential decay to the minima of sliding 6.8 s windows. The trace was then divided by the fit. ΔF/F was computed relative to baseline fluorescence calculated as the mean of the first 2 min of recording prior to sound presentation.

For sound-aligned analyses, fluorescence traces were segmented into trials time-locked to CS onset and averaged within each mouse. For shock-aligned analyses, traces were aligned to foot shock onset during conditioning. For freezing-aligned analyses, ΔF/F traces were aligned to freezing onset and offset, defined from the freezing annotation.

Peak response amplitude was defined as the maximum ΔF/F value within a predefined post-stimulus window (0-29 s after CS onset for sound-evoked responses; 0-10 s after shock onset for shock-evoked responses). Latency to peak was defined as the time elapsed between stimulus onset and the time point of maximal ΔF/F within the corresponding analysis window. Peak amplitudes and latencies were first computed for each mouse based on the trial-averaged trace and then averaged across mice.

For testing attenuation or amplification to sounds, sound-evoked responses were averaged within each mouse across early (trials 1-5), middle (trials 6-10) and late (trials 11-15) trial blocks. Peak amplitudes were extracted from the trial-averaged trace for each block, resulting in one value per mouse and condition and were compared using one-way repeated-measures ANOVA.

To test whether shock responses were correlated with shock-induced locomotion, maximum ΔF/F and maximum speed within 0-10 s after shock were correlated, and linear regression, Pearson correlation and cross-correlation analyses were performed using custom-written MATLAB scripts.

For neuron–astrocyte comparisons, latency-to-peak values were statistically compared. In addition, freezing onset–related changes in ΔF/F were quantified by comparing mean ΔF/F values during the 0.15 s preceding freezing onset with at 2 s following freezing onset in neurons and astrocytes.

#### Effects of astrocytic Gi-GPCR activation on astrocytic Ca^2+^ signaling

For experiments assessing the effect of astrocytic Gi-GPCR activation on sound-evoked astrocytic Ca²⁺ responses during retrieval, mice expressing AAV-hGFAP-GCaMP6f and AAV-hGFAP-hM4D_i_ in TeA were tested in a within-session pre/post design. Two CS⁺ and two CS⁻ presentations were delivered before i.p. injection of CNO or saline, followed by two additional CS⁺ and two CS⁻ presentations 30 min after injection. Animals remained in the conditioning boxes throughout the entire session and fluorescence signals were recorded continuously.

ΔF/F was computed for the entire trace, with F_0_ defined as the mean fluorescence during the first 2 min of the session, thereby preserving potential baseline shifts induced by hM4D_i_ activation.

For visualization of CS^+^-evoked activity, responses were first averaged across trials within each animal for pre- and post-injection periods. These per-animal averages were then averaged across animals to obtain group mean traces.

For peak response quantification, each trial was baseline corrected by subtracting the mean ΔF/F during the pre-stimulus window. Peaks were defined as the maximum ΔF/F within the CS^+^ time window. Baseline astrocytic activity was quantified as the mean ΔF/F during 2.4 s preceding CS⁺ presentations before and after i.p. injections of saline or CNO. Changes in both peak responses and baseline activity were calculated as POST-PRE differences and statistically compared.

#### Projection-specific test of the effect of astrocytic Gi-GPCR activation

To assess whether acute activation of astrocytic Gi-GPCR signaling alters spontaneous neuronal output from the TeA to the amygdala/posterior striatum, we injected AAV-CBA-DIO–GCaMP6s and AAV-hGFAP-hM4D_i_ into the TeA, and AAV2-retro-hSyn1-chl-iCre with AAV-hSyn1-DIO-mCherry into the lateral amygdala/posterior striatum. Mice had an optical fiber implanted dorsal to TeA. First, mice underwent the same auditory fear conditioning paradigm as described above with an i.p. injection of CNO (4 mg/kg^-1^) or saline delivered on the retrieval day. On one of the following days, an additional experiment was performed in which an IP injection of CNO or saline was administered immediately before the start of the session. Mice were placed in the behavioral box with the shock grid covered by an acrylic plate and no auditory stimuli were presented. Fluorescence activity was recorded continuously for 60 min. Signal processing and ΔF/F calculations were performed as described above.

Synchronized calcium events were detected after light denoising of ΔF/F traces using a moving median filter (0.1 s window), followed by application of an adaptive threshold defined as the local mean plus two local standard deviations computed over a 500 s sliding window. Peaks exceeding this threshold and separated by at least 0.4 s were classified as calcium events. A continuous event rate was obtained by convolving the binary spike train with a 60 s window. To quantify synchronized activity, the mean event rate was calculated over the final 30 min of the recording period for the final recording day.

#### Vaginal smear

Vaginal smears were collected from 20 C57BL/6JRj control mice 30 min after retrieval, following established procedures by McLean et al. (2012). Samples were stained with hematoxylin solution (Sigma, cat. no. HHS16-500ML) and examined under a light microscope. Estrus stages were classified independently by two experimenters, and CS^+^ mean freezing levels were compared across stages.

#### Exclusion criteria

Mice were excluded if histology revealed absent viral expression or misplaced injection sites, defined as injections not centered in the TeA, with the majority of expression located outside of TeA, or showing spread into non-cortical adjacent regions (e.g., hippocampal formation). Additional exclusion criteria included technical failures during behavior or insufficient fluorescence (baseline fluorescence <3x autofluorescence for fiber-implanted mice).

### Auditory go/no-go discrimination task

Female mice (*N* = 20-24 per experiment, 8-9 weeks old) received bilateral TeA injections of either AAV-hSyn1-dlox-hM4D_i_ co-injected with AAV-mCaMKIIα-iCre or AAV-hGFAP-hM4D_i_ as described above. To assess whether CNO alone altered behavior, an additional cohort of female C57BL/6JRj mice (*N* = 20) underwent the same behavioral paradigm without AAV injection.

The behavioral boxes were custom-built from acrylic plates (30 x 15.5 x 30 cm) and equipped with infrared LED illumination, a Titanium Dome Tweeter (TW025A20 1, Audax) delivering amplified sounds (55 dBA at mouse height) from a SparkFun MP3-player shield (12660. SparkFun Electronics), and a Blackfly S USB3 camera (BFS-U3-13Y3M-C, Teledyne) recording at 30 fps. One wall contained a nose-port connected via tubing to an Arduino Uno-controlled rotating pellet dispenser. An infrared beam was used to detect nose pokes. An Arduino Uno board was programmed to trigger the pellet dispenser whenever a beam break occurred during a CS^+^ presentation. All Arduino TTL outputs (LEDs, MP3 player, pellet dispenser), and Arduino inputs (beam breaks) were acquired through a T7 Daq Board (LabJack) and monitored and analyzed online with a custom-written MATLAB script.

The behavioral timeline was minimized to avoid neurotoxicity from long-term AAV expression. Mice were handled beginning on day 5 after surgery and single-housed starting after day 2 of handling. After seven days of handling, mice were weighed on two consecutive days, and the mean value was defined as baseline body weight. Food restriction was then initiated: over 5-8 days, weight was gradually reduced to 90% of baseline and maintained at 80.5-85% thereafter, with daily reductions not exceeding 3% to minimize distress. After reaching <90% baseline weight, five sucrose pellets (Bio-Serv, cat. no. F07595) were placed into each home cage for taste familiarization. Training began the following day.

The training protocol was adapted from Horst et al. (2012), with one session per day conducted during the dark phase at ZT14-19. Each session terminated after retrieval of 20 pellets or after a maximum duration (20 min for early stages, 30 min for later stages and the experimental phase). Mice were trained for 6-24 days before entering the experimental phase.

Training proceeded through the following stages:

Step 1: Nose pokes were rewarded with pellet delivery (20-26 s inter-reward interval). Advancement required consumption of >10 pellets.

Step 2: A 5 s, 8.37 kHz (the perceptual/logarithmic midpoint of the 7 and 10 kHz sounds used for the final experiments) training tone was introduced every 15-21 s. Nose pokes during the tone were rewarded, whereas poking during inter-tone intervals triggered a 4.5-13 s delay. Advancement required consumption of >10 pellets.

Step 3: The delay for poking between tones increased to 9-18 s, advancement required consumption of >15 pellets.

Step 4: The task structure resembled the final experimental phase: a silent “catch” trial occurred in 50% of trials. Either the 8.37 kHz sound or the silent trial occurred every 10-16 s. Poking during inter-trial intervals triggered a 9-18 s delay. To advance, mice had to consume >15 pellets, complete the session within 30 min, and achieve a hit rate >50%.

Step 5: An additional 15 s penalty was applied for poking during silent false-alarm (FA) trials. Advancement required consumption of >15 pellets, a hit rate >50%, and a discrimination index (DI) >0.5.

During the experimental phase, two 5 s pure tones were presented: 7 kHz and 10 kHz. One tone served as the CS^+^, for which nose poking during the sound resulted in pellet delivery (hit), and the other as the CS^-^, for which poking triggered a 15 s delay (FA). Tone identity (CS^+^/CS^-^) was counterbalanced across groups. Trials occurred every 10-16 s, and poking during inter-trial intervals produced a 9-18 s delay. Mice were subjected to the experimental phase for 21 days. For days 2-6, mice received an i.p. injection of CNO (4 mg/kg^-1^) or saline 30 min before testing.

#### Behavioral analysis

Four behavioral measures were quantified: the hit rate (HR), false alarm rate (FAR), discrimination index (DI), and discriminability (D’). HR was calculated as hits/(hits+misses). FAR was calculated as FA/(FA+CR), where FA is the number of times when the mouse incorrectly pokes during the presentation of the CS^-^ and correct rejections (CR) are the number of times when the mouse correctly avoids poking during the presentation of the CS^-^. The DI was calculated as (HR-FAR)/(HR+FAR), assessing the degree of stimulus discrimination. D’ was computed as the difference between the inverse cumulative normal of HR and FAR.

In addition, latencies to respond to CS^+^ and baseline magazine entries were quantified. Baseline magazine entries were calculated as the average probability of a mouse entering the magazine in no-sound periods. A 5-s analysis window starting 5 s after each unpaired sound was used. This period was chosen due to its low likelihood of being affected by false alarm entries, hit entries and sugar pellet consumption entries.

To compare groups statistically, performance metrics were calculated for each session and mouse, then averaged over the days of CNO/saline i.p. injections (sessions 2-6 in the experimental phase).

### Quantification and statistical analysis

Group sizes are indicated in figure legends; power calculations were used to predetermine sample sizes.

All analyses were performed in MATLAB using custom-written scripts unless indicated otherwise. Data are presented as mean ± SEM or median [IQR]. All statistical tests were two-sided. For parametric tests, assumptions of normality and homogeneity of variance were verified using Shapiro-Wilk and *F*-tests. When these assumptions were violated, Mann-Whitney *U* tests or Kruskal-Wallis tests were used. Otherwise, paired or unpaired *t*-tests were applied. For comparisons involving repeated measurements within the same subjects, one-way repeated-measures ANOVA was used. Where multiple related hypotheses were tested, *p*-values were adjusted for multiple comparisons using the Bonferroni-Holm method with family-wise α = 0.05. For statistical comparisons of freezing data, *p*-values were corrected for testing of three related measures: baseline freezing, CS^+^ freezing, and discrimination index.

For auditory fear conditioning experiments, behavioral readouts were analyzed on a per-mouse basis and compared between experimental and control groups.

#### Blinding strategy

For all behavioral experiments, experimenters were blinded to group assignment throughout the entire experimental workflow, including surgeries, behavioral testing, analysis, histological validation and exclusion of mice. Mice were randomly assigned to groups. Groups that were statistically compared in this study were run during the same time period and, wherever possible, in parallel.

## Data availability

Data reported in this paper will be shared by the lead contact upon request. Any additional information required to reanalyze the data reported in this paper is available from the lead contact upon request.

## Code availability

All original code is available from the lead contact upon request.

## Acknowledgements

We thank Philipp Velicky and the team of the Imaging Core Facility for technical support. We thank Bryan Roth and Yulong Li for providing Addgene plasmids. We thank the animal care staff (Sonja Reynoso de Leon, Aurica Jelinek, Corina Brottrager, Christian Schönauer and Jutta Pilecky) for their support. We thank Hugo Malagon-Vina for assistance with data analysis. We thank Greta Grudeli for feedback on the manuscript.

This work was funded by the Vienna Science and Technology Fund (WWTF) and the City of Vienna through project VRG21-015. L.F., Y.A.S., M.H., and A.C.-M. were supported by NIH R01 (NIMH, 7R01MH129732-04, A.C.-M.), CU Anschutz startup funding (A.C.-M.), NIH RF1NS128739 (M.H.), and CU Anschutz SOM ASPIRE Program (M.H.).

## Author contributions

Conceptualization: S.N.H., S.M.. Data analysis: S.N.H., S.M., A.C.M., V.S., R.R., L.K., Y.A.S.. Investigation: S.N.H., A.C.M., V.S., L.F., R.R.. Writing: S.N.H., S.M., V.S., A.C.M., A.C-M.. Visualization: S.N.H., S.M., A.C.M, V.S.. Supervision: S.M., A.C.-M., M.H.. Funding acquisition: S.M., A.C.-M., M.H.. Ethics declaration Competing interests: None to be declared.

## Extended Data

**Extended Data Figure 1.**
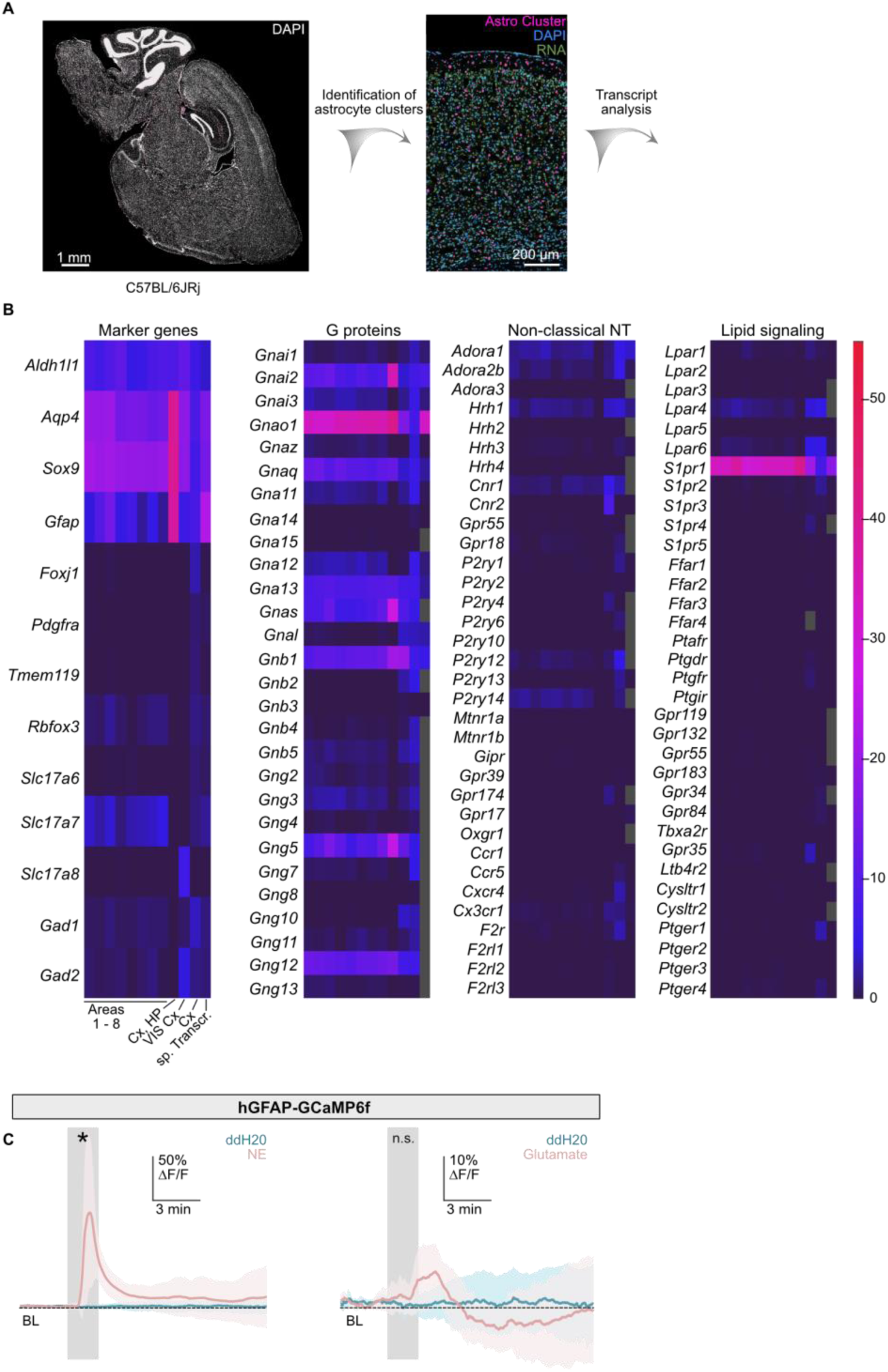
(a) Left, exemplary sagittal section with DAPI staining used for spatial transcriptomics analysis. Right, higher-magnification image of cortical area showing ROIs belonging to the astrocyte-rich cluster. (b) Heatmaps showing normalized expression of selected marker genes, G protein subunits and GPCRs across transcriptomic datasets. Columns correspond to Sugino et al. (2019), Tasic et al. (2016), Yao et al. (2021), single-cell RNA-seq datasets and Xenium spatial transcriptomics data, as indicated in Fig. 1b. Grey fields indicate data not available. (c) Astrocytic Ca²⁺ imaging in acute TeA slices expressing AAV-hGFAP-GCaMP6f. Ligand application is indicated by the grey shaded area. Peak signals: norepinephrine (NE) 90.46% ± 8.09% ΔF/F; glutamate 9.75% ± 0.89% ΔF/F; ddH_2_O 3.4% ± 0.59%. Significant increases in GCaMP6f fluorescence are indicated by asterisks (NE: *p* ≤ 0.01; glutamate: *p* = 0.073; unpaired *t*-test, *N* = 6 slices per condition).

**Extended Data Figure 2.**
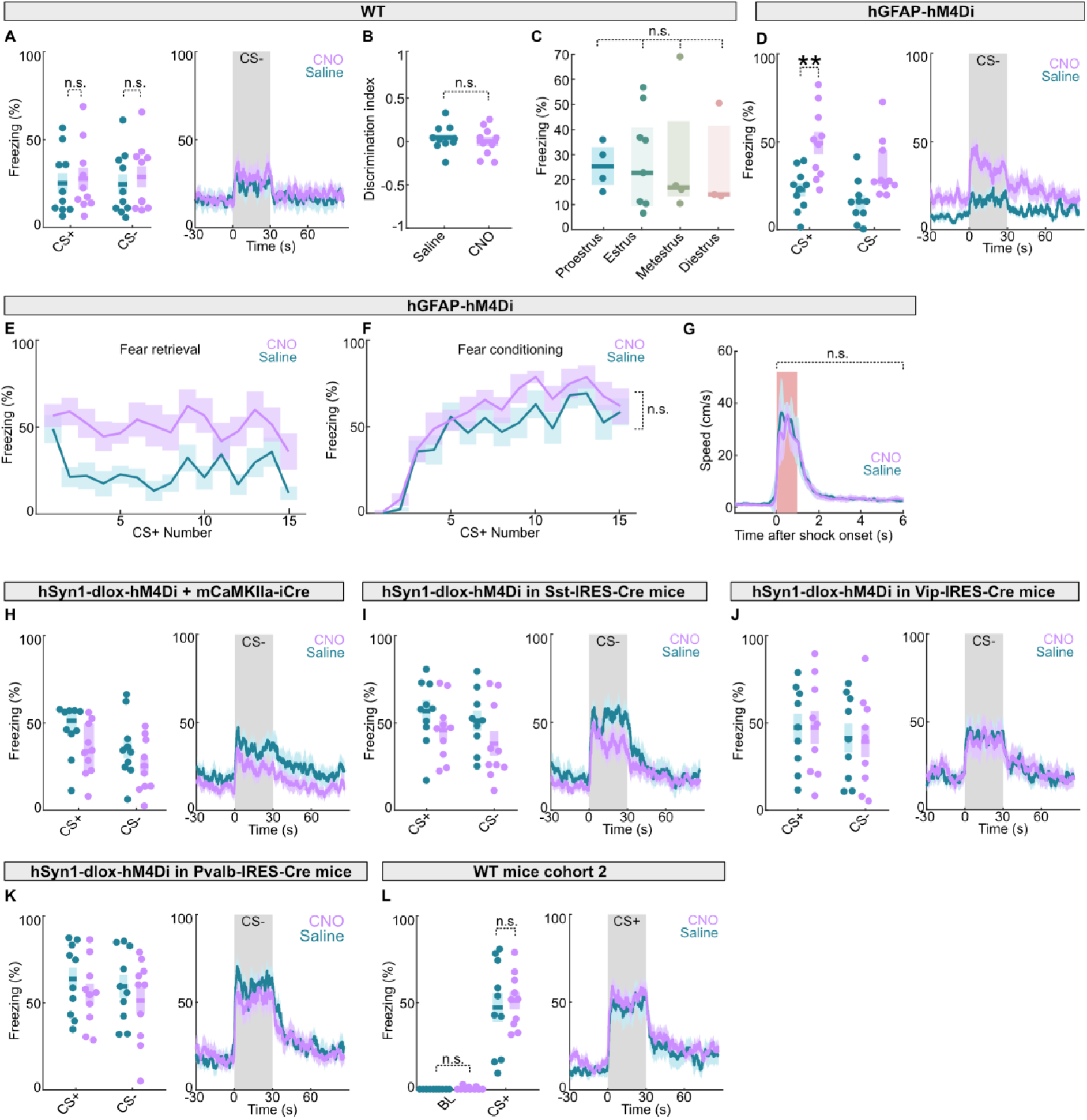
(a) Quantification and time course of freezing levels to CS^+^ and CS^−^ as indicated during retrieval in naïve wild-type mice (CS^-^ CNO: 28.86 ± 6.03%, saline: 24.60 ± 5.75%). CNO did not significantly change freezing upon CS^+^ or CS^-^ presentation (CS^-^ *p* = 0.616, unpaired *t-*test), *N* = 10 mice per condition. (b) Discrimination index for naïve wild-type mice treated with saline or CNO during retrieval. (CNO: – 0.017 ± 0.052; saline: 0.024 ± 0.046; *p* = 1, unpaired *t*-test), *N* = 10 per condition. (c) Freezing levels upon CS⁺ presentation during retrieval as a function of estrous stage (proestrus (25.25% [17.88-32.98%], *n* = 4), estrus (22.71% [9.5-40.83%], *n* = 9), metestrus (16.87% [13.33-43.34%], *n* = 4), diestrus (14.11% [13.63-41.44%], *n* = 3); *p* = 0.95, Kruskal-Wallis test), *N* = 20. (d) Same as in (a), but for mice expressing AAV-hGFAP-hM4D_i_-mCherry in TeA astrocytes, *N* = 10 per condition. (e) Time course of freezing levels upon presentation of 15 CS^+^ during retrieval for mice expressing AAV-hGFAP-hM4D_i_-mCherry in TeA astrocytes, treated with saline or CNO, *N* = 10 per condition. (f) Time course of freezing levels upon presentation of 15 CS⁺ during conditioning for mice receiving saline or CNO treatment on the following day. There was no difference in freezing to the last CS^+^ (CNO: 61.43 ± 3.25%; saline: 58.11 ± 2.45%; *p* = 0.8, unpaired *t*-test), *N* = 10 per condition. (g) Mean shock-evoked locomotor response during fear conditioning in AAV-hGFAP-hM4D_i_-mCherry-expressing mice assigned to saline or CNO treatment. Shock period is indicated by the shaded region (Area under the curve (AUC): CNO: 52.35 ± 2.04; saline: 55.21 ± 3.21; *p* = 0.463, unpaired *t*-test), *N* = 10 per condition. (h) Same as in (a), but for AAV-hSyn1-dlox-hM4D_i_-mCherry + mCaMKIIα-iCre expression in TeA, *N* = 10 per condition. (i) Same as in (a), but for Sst-IRES-Cre mice expressing AAV-hSyn1-dlox-hM4D_i_-mCherry in TeA, N = 10 per condition. (j) Same as in (a), but for Vip-IRES-Cre mice expressing AAV-hSyn1-dlox-hM4D_i_-mCherry in TeA, *N* = 9 per condition. (k) Same as in (a), but for Pvalb-IRES-Cre mice expressing AAV-hSyn1-dlox-hM4D_i_-mCherry in TeA, *N* = 10 per condition. (l) Control experiment to test off-target CNO effects in a second cohort of naïve wild-type mice. Left: baseline freezing (CNO: 0.00% [0.00-1.15%], saline: 0.00% [0.00-0.00%]; *p* = 0.093, Mann-Whitney *U* test) and CS⁺ (CNO: 51.22 ± 4.95%, saline: 47.38 ± 8.28%; *p* = 0.698, unpaired *t*-test) for saline- and CNO-treated animals. Right: peri-stimulus freezing time course aligned to CS⁺ onset (CS⁺ interval shaded), *N* = 10 per condition.

**Extended Data Figure 3.**
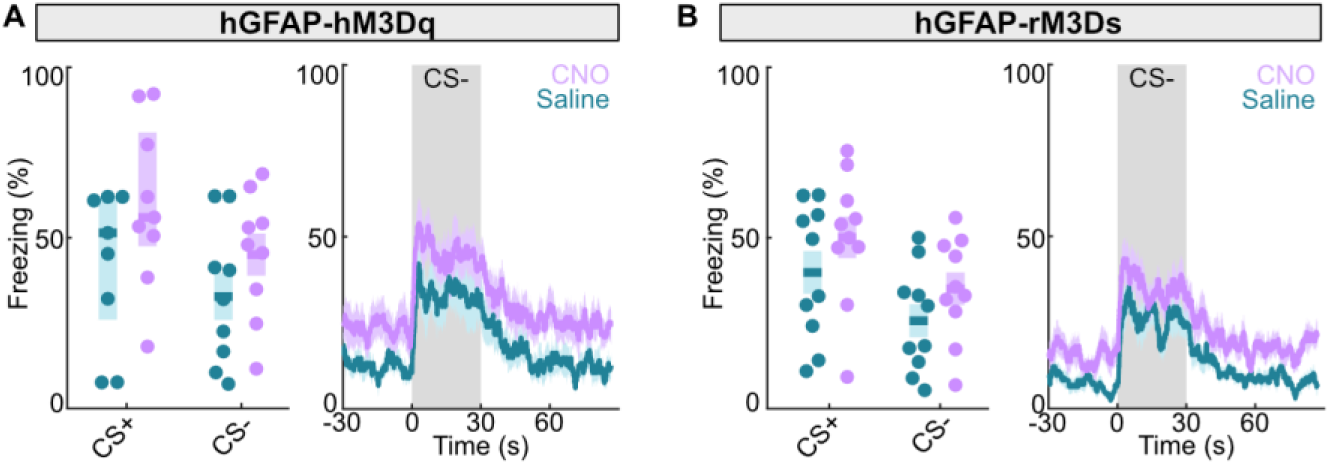
(a) Quantification and time course of freezing levels to CS^+^ and CS^−^ as indicated during retrieval in mice expressing AAV-hGFAP-hM3D_q_-mCherry in TeA, *N* = 9 mice per condition. (b) Same as in (a), but for mice expressing hGFAP-HA-rM3D_s_-mCitrine in TeA, *N* = 10 mice per condition.

**Extended Data Figure 4.**
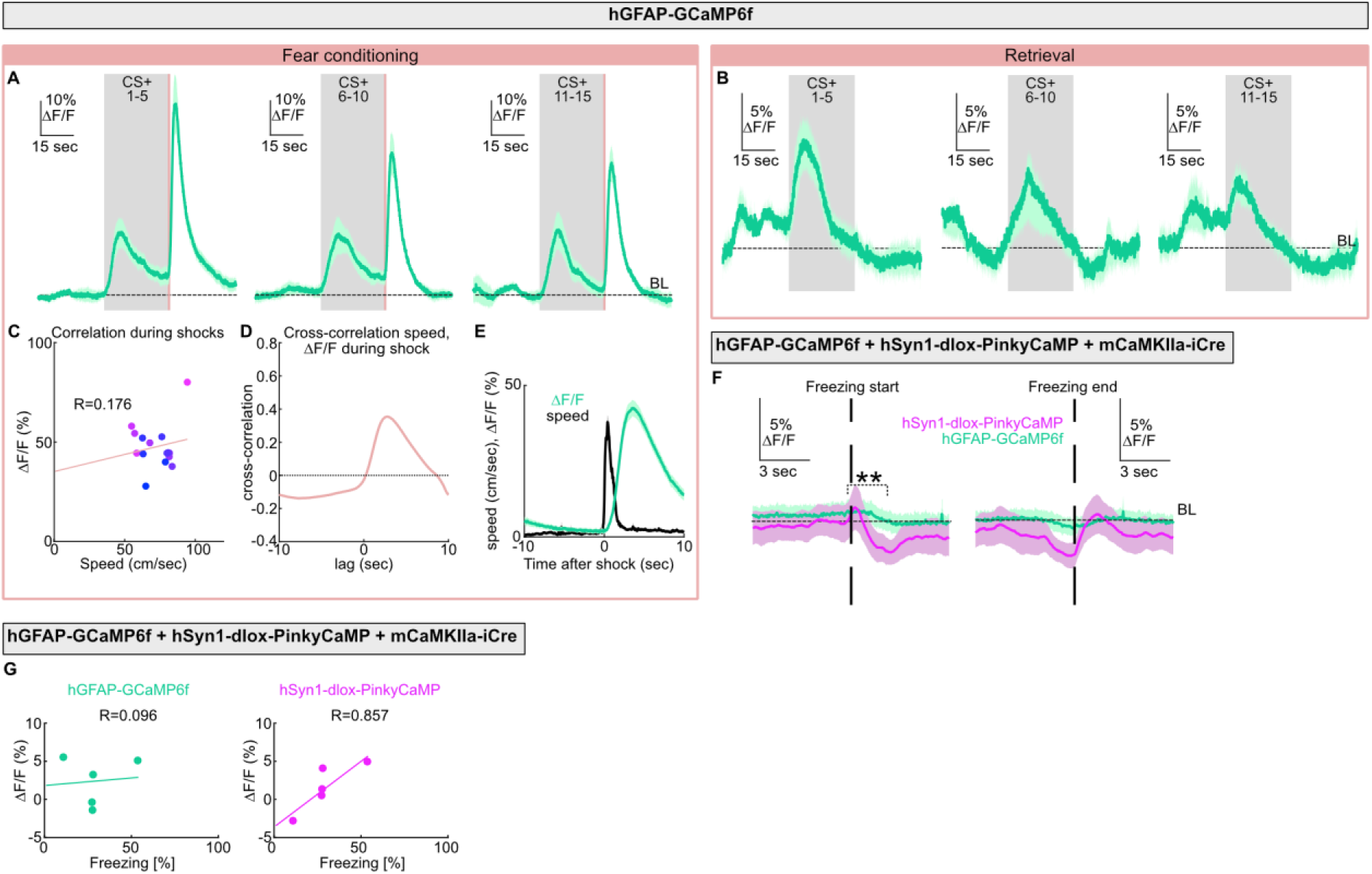
(a) Normalized fluorescence changes (ΔF/F) from AAV-hGFAP-GCaMP6f-expressing astrocytes aligned to CS⁺ presentations during fear conditioning. Traces are grouped by trial number (CS⁺ 1–5, 6–10, and 11–15). Peak ΔF/F CS⁺ 1-5: 20.9 ± 1.25%, peak ΔF/F CS^+^ 6–10: 18.84 ± 1.62%; peak ΔF/F CS^+^ 11-15: 19.19 ± 1.79%. No statistical difference in peak ΔF/F during CS^+^ presentations across trial blocks was detected (*p* = 0.866; one-way repeated-measures ANOVA), *N* = 6 mice. (b) Normalized fluorescence changes from AAV-hGFAP-GCaMP6f-expressing astrocytes aligned to CS⁺ presentations during retrieval, grouped by trial number (CS⁺ 1-5, 6-10, and 11-15). Peak ΔF/F CS⁺ 1-5: 9.83 ± 0.85%; peak ΔF/F CS⁺ 6-10: 7.24 ± 0.96%; peak ΔF/F CS⁺ 11-15: 5.66 ± 0.51%. No statistical difference in peak ΔF/F during CS^+^ presentations across trial blocks was detected (*p* = 0.237; one-way repeated-measures ANOVA), *N* = 6 mice. (c) Correlation between maximum locomotor speed and maximum astrocytic Ca²⁺ transients across all shocks (+10 s), averaged across mice. Each point represents the mean response to a given shock (1 (pink) to 15 (blue)). Linear regression and Pearson correlation coefficient are shown, *N* = 6 mice. (d) Cross-correlation between locomotor speed and astrocytic Ca²⁺ ΔF/F during the shock period (±10 s), averaged across mice, *N* = 6 mice. (e) Mean speed and astrocytic ΔF/F during the shock period (±10 s), averaged across mice, *N* = 6 mice. (f) Normalized fluorescence changes from AAV-hGFAP-GCaMP6f-expressing astrocytes and AAV-hSyn1-dlox-PinkyCaMP-expressing neurons aligned to freezing onset (left) and freezing offset (right). Change in ΔF/F from 0.15 s before to 2 s after freezing onset: astrocytes: mean ± SEM = –0.62 ± 0.26%; neurons: –2.54 ± 0.57% (statistical comparison between neurons and astrocytes: *p* = 0.025; unpaired *t*-test), *N* = 5 mice. (g) Quantification of correlation between average Ca^2+^ transients during CS^+^ and freezing level in astrocytes (left) and neurons (right). Dot plot with linear regression and Pearson correlation, *N* = 5 mice.

**Extended Data Figure 5.**
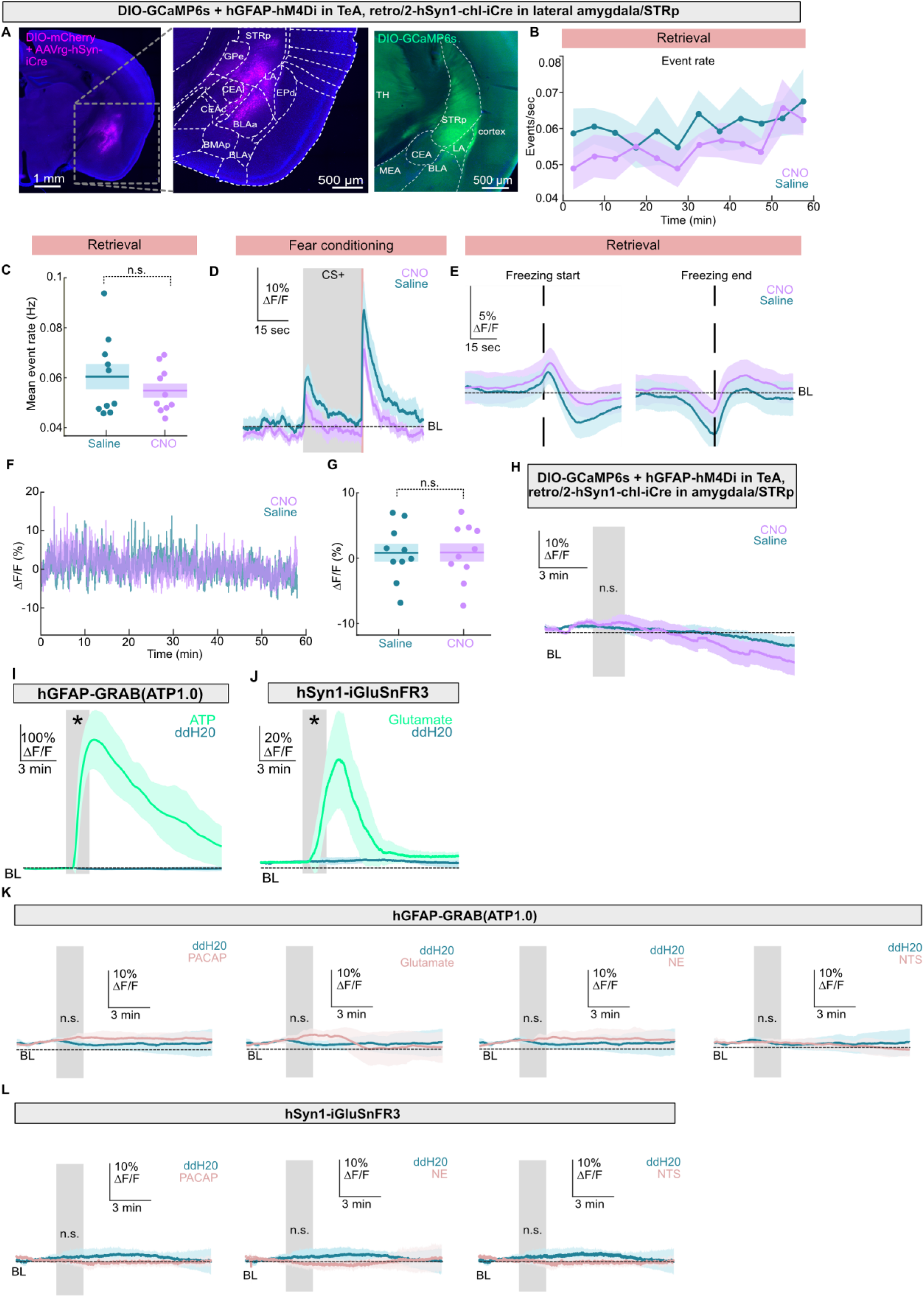
(a) Representative epifluorescence images showing mCherry-labeled injection site in STRp and amygdala after injection of AAV2-retro-hSyn1-iCre and AAV-hSyn1-DIO-mCherry (left) and axonal projections following AAV-CBA-DIO-GCaMP6s injection in TeA (right). (b) Normalized fluorescence changes (ΔF/F) from amygdala/STRp-projecting neurons expressing AAV-CBA-DIO-GCaMP6s during fear conditioning before astrocytic hM4D_i_ was activated (CNO) or control treatment (saline). Peak ΔF/F during CS^+^ for CNO: 6.04 ± 0.55%; saline: 8.74 ± 1.03%, *N* = 9 mice per condition. (c) Normalized fluorescence changes from amygdala/STRp-projecting neurons aligned to freezing onset (left) and freezing offset (right), during astrocytic Gi-GPCR activation (CNO) or control treatment, *N* = 9 mice per condition. (d) Event rate of synchronized Ca²⁺ transients in amygdala/STRp-projecting neurons during retrieval following i.p. injection of CNO or saline, *N* = 9 mice per condition. (e) Mean synchronized event rate during retrieval for CNO and saline-treated animals (CNO: 0.05 ± 0.01 Hz; saline: 0.06 ± 0.01 Hz, *p* = 0.351, unpaired *t*-test), *N* = 9 mice per condition. (f) Mean fluorescence changes from amygdala/STRp-projecting neurons following CNO or saline injection during retrieval. *N* = 9 mice per condition. (g) Quantification of mean fluorescence changes during retrieval following CNO or saline injection. Mean ΔF/F: CNO: 0.87 ± 1.39%; saline: 0.82 ± 1.35%, *p* = 0.968, unpaired *t*-test, *N* = 9 mice per condition. (h) Ca^2+^ transients from amygdala/STRp-projecting neurons expressing AAV-CBA-DIO-GCaMP6s in acute mouse brain slices during bath application of CNO/saline (peak ΔF/F CNO: 3.17 ± 1.46%; saline: 5.62 ± 2.53%; *p* = 0.42; unpaired *t*-test). The grey shaded region indicates ligand application, *N* = 6 slices per condition. (i) Astrocytic ATP-evoked transients in acute TeA slices expressing AAV-hGFAP-GRAB(ATP1.0). Bath application of ATP (peak ΔF/F ATP: 366.8 ± 13.22%, ddH_2_O: 5.14 ± 0.30%; *p* ≤ 0.01; unpaired *t*-test), *N* = 6 slices per condition. (j) Glutamate-evoked transients in acute TeA slices expressing AAV-hSyn1-iGluSnFR3. Bath application of glutamate (peak ΔF/F glutamate: 72.45% [33.42-73.97%], ddH_2_O: 1.72% [1.47-3.72%]; *p* ≤ 0.01; Mann-Whitney *U* test), *N* = 6 slices per condition. (k) Astrocytic ligand-evoked transients in acute TeA slices expressing AAV-hGFAP-GRAB(ATP1.0). Bath application of PACAP (peak ΔF/F 7.64% ± 0.39%; *p* = 0.19), glutamate (9.41 ± 0.54%; *p* = 0.075), norepinephrine (NE; 7.39 ± 0.68%; *p* = 0.49), neurotensin (NTS; 5.06% ± 0.29%; *p* = 0.939) did not significantly alter ATP signals compared with ddH_2_O controls (peak ΔF/F 5.14% ± 0.30%; unpaired *t*-tests), *N* = 6 slices per condition. (l) Ligand-evoked transients in acute TeA slices expressing AAV-hSyn1-iGluSnFR3. Bath application of PACAP (peak ΔF/F median [IQR]; 1.83% [1.61-1.97%]; *p* = 1; Mann-Whitney *U* test), NE (0.62% ± 0.14%, *p* = 0.158; unpaired *t*-test), or NTS (peak ΔF/F 0.7% ± 0.13%, *p* = 0.158; unpaired *t*-test), did not alter glutamate signals compared with ddH_2_O controls (parametric estimate 2.25% ± 0.27%; non-parametric estimate 1.72% [1.47-3.72%]), *N* = 6 slices per condition. Abbreviations: BLAa, basolateral anterior amygdalar nucleus, anterior; BMAp, basomedial amygdala, posterior; BMAv, basomedial amygdala, ventral; CEAc, central amygdala, central; CEAl, central amygdala, lateral; EPd, dorsal endopiriform nucleus; GPe, external globus pallidus; LA, lateral amygdala; MEA, medial amygdala; STRp, posterior striatum; TH, thalamus

**Extended Data Figure 6.**
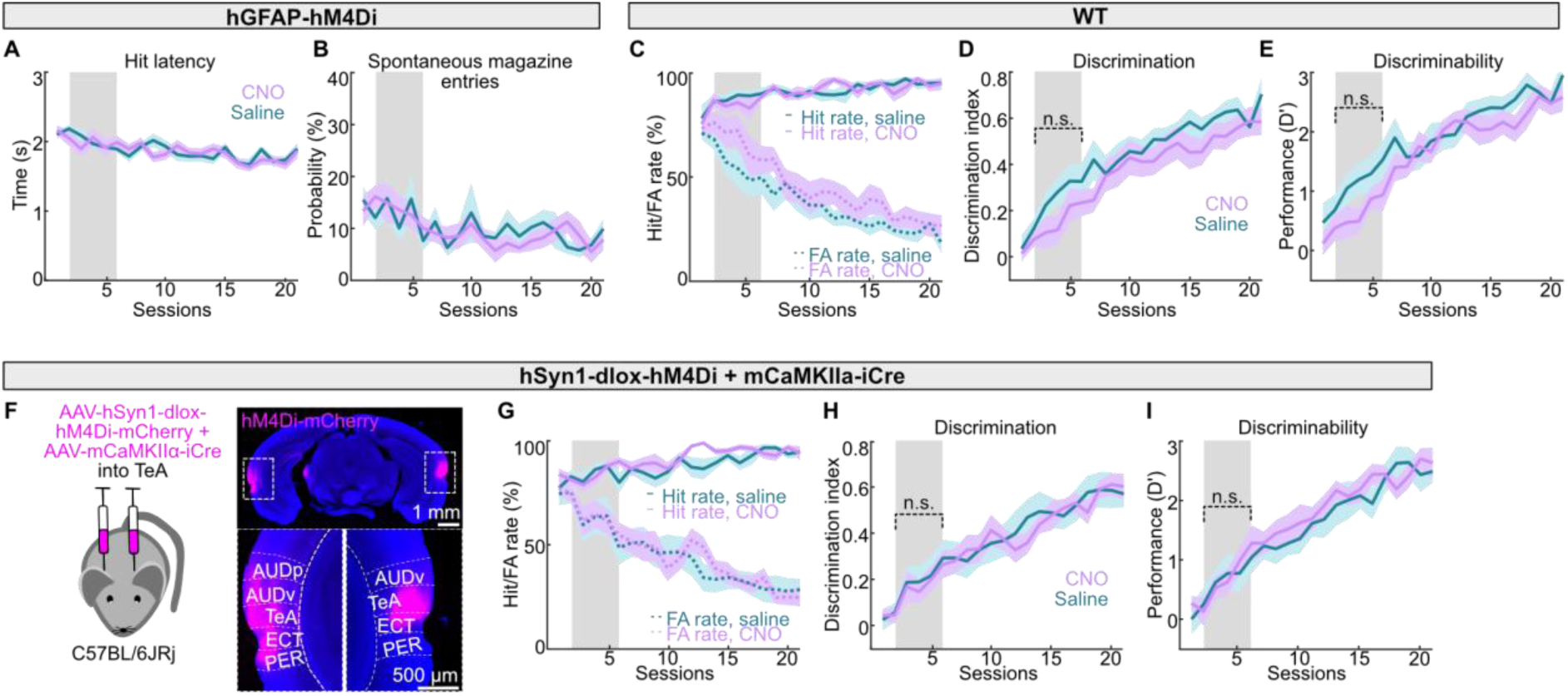
(a) Latency of hits across sessions in CNO- and saline-treated mice expressing AAV-hGFAP-hM4D_i_-mCherry, *N* = 11 mice per condition. (b) Probability of magazine entries in no-sound periods across sessions in CNO- and saline-treated mice expressing AAV-hGFAP-hM4D_i_-mCherry, *N* = 11 mice per condition. (c) Hit rate and false alarm (FA) rate in CNO- and saline-treated naïve mice, *N* = 10 mice per condition. (d) Discrimination index of naïve mice (mean across sessions 2-6; CNO: 0.15 ± 0.05; saline: 0.26 ± 0.06, *p* = 0.199, unpaired *t*-test), *N* = 10 mice per condition. (e) Discriminability (D’) of naïve mice (mean across sessions 2-6; CNO: 0.63 ± 0.06; saline: 1.15 ± 0.07, *p* = 0.199, unpaired *t*-test), *N* = 10 mice per condition. (f) Exemplary epifluorescence image showing expression of AAV-hSyn1-dlox-hM4D_i_-mCherry combined with AAV-mCaMKIIα-iCre. (g) Hit rate and FA rate across sessions in mice with hM4D_i_ expression in TeA mCaMKIIα^+^ neurons, treated with CNO or saline, *N* = 12 mice per condition. (h) Discrimination index across sessions in mice with hM4D_i_ expression in TeA mCaMKIIα^+^ neurons, treated with CNO or saline (median [IQR] across sessions 2-6; CNO: 0.11 [0.02-0.39]; saline: 0.08 [0.01-0.39] *p* = 1, Mann-Whitney *U* test), *N* = 12 mice per condition. (i) Discriminability (D’) across sessions in mice with hM4D_i_ expression in TeA mCaMKIIα^+^ neurons, treated with CNO or saline (mean across sessions 2-6; CNO: 0.71 ± 0.06; saline: 0.69 ± 0.07, *p* = 1, unpaired *t*-test), *N* = 12 mice per condition. Abbreviations: AUDv, ventral auditory area; AUDp, primary auditory area; ECT, ectorhinal area; HPF, hippocampal formation; PER, perirhinal area; TeA, temporal association area

## References

1. Adamsky, A., Kol, A., Kreisel, T., Doron, A., Ozeri-Engelhard, N., Melcer, T., Refaeli, R., Horn, H., Regev, L., Groysman, M., London, M., & Goshen, I. (2018). Astrocytic Activation Generates De Novo Neuronal Potentiation and Memory Enhancement. Cell, 174(1), 59–71.e14. 10.1016/j.cell.2018.05.002

2. Allen Institute for Brain Science. (2011). Allen Mouse Brain Atlas [dataset]. Available from Mouse.Brain-Map.Org.

3. Araque, A., Carmignoto, G., Haydon, P. G., Oliet, S. H. R., Robitaille, R., & Volterra, A. (2014). Gliotransmitters Travel in Time and Space. Neuron, 81(4), 728–739. 10.1016/j.neuron.2014.02.007

4. Bedwell, S. A., Billett, E. E., Crofts, J. J., MacDonald, D. M., & Tinsley, C. J. (2015). The topology of connections between rat prefrontal and temporal cortices. Frontiers in Systems Neuroscience, 9. 10.3389/fnsys.2015.00080

5. Bekar, L. K., He, W., & Nedergaard, M. (2008). Locus Coeruleus α-Adrenergic–Mediated Activation of Cortical Astrocytes In Vivo. Cerebral Cortex, 18(12), 2789–2795. 10.1093/cercor/bhn040

6. Bergles, D. E., & Jahr, C. E. (1997). Synaptic Activation of Glutamate Transporters in Hippocampal Astrocytes. Neuron, 19(6), 1297–1308. 10.1016/S0896-6273(00)80420-1

7. Bettler, B., Kaupmann, K., Mosbacher, J., & Gassmann, M. (2004). Molecular Structure and Physiological Functions of GABA _B_ Receptors. Physiological Reviews, 84(3), 835–867. 10.1152/physrev.00036.2003

8. Bukalo, O., O’Sullivan, R., Tanisumi, Y., Mendez, A., Weinholtz, C., Zimmerman, S., Offenberg, V., Carpenter, O., Bhagwat, H., Mosley, S., O’Malley, J. J., Lyons, K., Fang, Y., Goldschlager, J., Ostroff, L. E., Penzo, M. A., Wake, H., Halladay, L. R., & Holmes, A. (2026). Astrocytes enable amygdala neural representations supporting memory. Nature. 10.1038/s41586-025-10068-0

9. Chai, H., Diaz-Castro, B., Shigetomi, E., Monte, E., Octeau, J. C., Yu, X., Cohn, W., Rajendran, P. S., Vondriska, T. M., Whitelegge, J. P., Coppola, G., & Khakh, B. S. (2017). Neural Circuit-Specialized Astrocytes: Transcriptomic, Proteomic, Morphological, and Functional Evidence. Neuron, 95(3), 531–549.e9. 10.1016/j.neuron.2017.06.029

10. Cheng, R., Zhong, W., Yan, Y., Yao, L., Yin, P., Xu, Z., Qin, X., Tan, J., Zeng, Y., Liu, J., & Xiao, Z. (2025). A disinhibitory microcircuit in the temporal association cortex for fear retrieval to pure tones. Scientific Reports, 15(1), 20457. 10.1038/s41598-025-05566-0

11. D Edelstein, A., A Tsuchida, M., Amodaj, N., Pinkard, H., D Vale, R., & Stuurman, N. (2014). Advanced methods of microscope control using μManager software. Journal of Biological Methods, 1(2), 1. 10.14440/jbm.2014.36

12. Dalmay, T., Abs, E., Poorthuis, R. B., Hartung, J., Pu, D.-L., Onasch, S., Lozano, Y. R., Signoret-Genest, J., Tovote, P., Gjorgjieva, J., & Letzkus, J. J. (2019). A Critical Role for Neocortical Processing of Threat Memory. Neuron, 104(6), 1180–1194.e7. 10.1016/j.neuron.2019.09.025

13. Drummond, G. T., Natesan, A., Celotto, M., Shih, J., Ojha, P., Osako, Y., Park, J., Sipe, G. O., Jenks, K. R., Breton-Provencher, V., Simpson, P. C., Panzeri, S., & Sur, M. (2024). Cortical norepinephrine-astrocyte signaling critically mediates learned behavior. 10.1101/2024.10.24.620009

14. Durkee, C. A., Covelo, A., Lines, J., Kofuji, P., Aguilar, J., & Araque, A. (2019). G _i/o_ protein-coupled receptors inhibit neurons but activate astrocytes and stimulate gliotransmission. Glia, 67(6), 1076–1093. 10.1002/glia.23589

15. Feigin, L., Tasaka, G., Maor, I., & Mizrahi, A. (2021). Sparse Coding in Temporal Association Cortex Improves Complex Sound Discriminability. The Journal of Neuroscience, 41(33), 7048–7064. 10.1523/JNEUROSCI.3167-20.2021

16. Fink, R., Imai, S., Gockel, N., Lauer, G., Renken, K., Wietek, J., Lamothe-Molina, P. J., Fuhrmann, F., Mittag, M., Ziebarth, T., Canziani, A., Kubitschke, M., Kistmacher, V., Kretschmer, A., Sebastian, E., Schmitz, D., Terai, T., Gründemann, J., Hassan, S., … Masseck, O. A. (2024). PinkyCaMP a mScarlet-based calcium sensor with exceptional brightness, photostability, and multiplexing capabilities. 10.1101/2024.12.16.628673

17. Ghenissa, O., Guayasamin, M., Ngo, K., Duquenne, M., Peyrard, S., Amilhon, B., & Murphy-Royal, C. (2026). Basolateral amygdala astrocytes encode anxiety states. Neuron. 10.1016/j.neuron.2026.02.038

18. González-Arias, C., Sánchez-Ruiz, A., Esparza, J., Sánchez-Puelles, C., Arancibia, L., Ramírez-Franco, J., Gobbo, D., Kirchhoff, F., & Perea, G. (2023). Dysfunctional serotonergic neuron-astrocyte signaling in depressive-like states. Molecular Psychiatry, 28(9), 3856–3873. 10.1038/s41380-023-02269-8

19. Gungor Aydin, A., Lemenze, A., & Bieszczad, K. M. (2024). Functional diversities within neurons and astrocytes in the adult rat auditory cortex revealed by single-nucleus RNA sequencing. Scientific Reports, 14(1), 25314. 10.1038/s41598-024-74732-7

20. Guttenplan, K. A., Maxwell, I., Santos, E., Borchardt, L. A., Manzo, E., Abalde-Atristain, L., Kim, R. D., & Freeman, M. R. (2025). GPCR signaling gates astrocyte responsiveness to neurotransmitters and control of neuronal activity. Science, 388(6748), 763–768. 10.1126/science.adq5729

21. Horst, N. K., Heath, C. J., Neugebauer, N. M., Kimchi, E. Y., Laubach, M., & Picciotto, M. R. (2012). Impaired auditory discrimination learning following perinatal nicotine exposure or β2 nicotinic acetylcholine receptor subunit deletion. Behavioural Brain Research, 231(1), 170–180. 10.1016/j.bbr.2012.03.002

22. Horstmeyer, A., Cramer, H., Sauer, T., Müller-Esterl, W., & Schroeder, C. (1996). Palmitoylation of Endothelin Receptor A. Journal of Biological Chemistry, 271(34), 20811–20819. 10.1074/jbc.271.34.20811

23. Inoue, A., Raimondi, F., Kadji, F. M. N., Singh, G., Kishi, T., Uwamizu, A., Ono, Y., Shinjo, Y., Ishida, S., Arang, N., Kawakami, K., Gutkind, J. S., Aoki, J., & Russell, R. B. (2019). Illuminating G-Protein-Coupling Selectivity of GPCRs. Cell, 177(7), 1933–1947.e25. 10.1016/j.cell.2019.04.044

24. Janesick, A., Shelansky, R., Gottscho, A. D., Wagner, F., Williams, S. R., Rouault, M., Beliakoff, G., Morrison, C. A., Oliveira, M. F., Sicherman, J. T., Kohlway, A., Abousoud, J., Drennon, T. Y., Mohabbat, S. H., & Taylor, S. E. B. (2023). High resolution mapping of the tumor microenvironment using integrated single-cell, spatial and in situ analysis. Nature Communications, 14(1), 8353. 10.1038/s41467-023-43458-x

25. Kambe, Y., Yamauchi, Y., Thanh Nguyen, T., Thi Nguyen, T., Ago, Y., Shintani, N., Hashimoto, H., Yoshitake, S., Yoshitake, T., Kehr, J., Kawamura, N., Katsuura, G., Kurihara, T., & Miyata, A. (2021). The pivotal role of pituitary adenylate cyclase-activating polypeptide for lactate production and secretion in astrocytes during fear memory. Pharmacological Reports, 73(4), 1109–1121. 10.1007/s43440-021-00222-6

26. Kol, A., Adamsky, A., Groysman, M., Kreisel, T., London, M., & Goshen, I. (2020). Astrocytes contribute to remote memory formation by modulating hippocampal–cortical communication during learning. Nature Neuroscience, 23(10), 1229–1239. 10.1038/s41593-020-0679-6

27. Krashes, M. J., Koda, S., Ye, C., Rogan, S. C., Adams, A. C., Cusher, D. S., Maratos-Flier, E., Roth, B. L., & Lowell, B. B. (2011). Rapid, reversible activation of AgRP neurons drives feeding behavior in mice. Journal of Clinical Investigation, 121(4), 1424–1428. 10.1172/JCI46229

28. LeDoux, J. (1998). Fear and the brain: where have we been, and where are we going? Biological Psychiatry, 44(12), 1229–1238. 10.1016/S0006-3223(98)00282-0

29. Letzkus, J. J., Wolff, S. B. E., Meyer, E. M. M., Tovote, P., Courtin, J., Herry, C., & Lüthi, A. (2011). A disinhibitory microcircuit for associative fear learning in the auditory cortex. Nature, 480(7377), 331–335. 10.1038/nature10674

30. Li, H., Zhao, Y., Dai, R., Geng, P., Weng, D., Wu, W., Yu, F., Lin, R., Wu, Z., Li, Y., & Luo, M. (2025). Astrocytes release ATP/ADP and glutamate in flashes via vesicular exocytosis. Molecular Psychiatry, 30(6), 2475–2489. 10.1038/s41380-024-02851-8

31. Li, Y., Li, L., Wu, J., Zhu, Z., Feng, X., Qin, L., Zhu, Y., Sun, L., Liu, Y., Qiu, Z., Duan, S., & Yu, Y.-Q. (2020). Activation of astrocytes in hippocampus decreases fear memory through adenosine A1 receptors. ELife, 9. 10.7554/eLife.57155

32. Ma, X., Chen, P., Wei, J., Zhang, J., Chen, C., Zhao, H., Ferguson, D., McGee, A. W., Dai, Z., & Qiu, S. (2024). Protocol for Xenium spatial transcriptomics studies using fixed frozen mouse brain sections. STAR Protocols, 5(4), 103420. 10.1016/j.xpro.2024.103420

33. Mather, M., Clewett, D., Sakaki, M., & Harley, C. W. (2016). Norepinephrine ignites local hotspots of neuronal excitation: How arousal amplifies selectivity in perception and memory. Behavioral and Brain Sciences, 39, e200. 10.1017/S0140525X15000667

34. McLean, A. C., Valenzuela, N., Fai, S., & Bennett, S. A. L. (2012). Performing Vaginal Lavage, Crystal Violet Staining, and Vaginal Cytological Evaluation for Mouse Estrous Cycle Staging Identification. Journal of Visualized Experiments, (67). 10.3791/4389

35. Melzer, S., Newmark, E. R., Mizuno, G. O., Hyun, M., Philson, A. C., Quiroli, E., Righetti, B., Gregory, M. R., Huang, K. W., Levasseur, J., Tian, L., & Sabatini, B. L. (2021). Bombesin-like peptide recruits disinhibitory cortical circuits and enhances fear memories. Cell, 184(22), 5622–5634.e25. 10.1016/j.cell.2021.09.013

36. Menegas, W., Akiti, K., Amo, R., Uchida, N., & Watabe-Uchida, M. (2018). Dopamine neurons projecting to the posterior striatum reinforce avoidance of threatening stimuli. Nature Neuroscience, 21(10), 1421–1430. 10.1038/s41593-018-0222-1

37. Nagai, J., Rajbhandari, A. K., Gangwani, M. R., Hachisuka, A., Coppola, G., Masmanidis, S. C., Fanselow, M. S., & Khakh, B. S. (2019). Hyperactivity with Disrupted Attention by Activation of an Astrocyte Synaptogenic Cue. Cell, 177(5), 1280–1292.e20. 10.1016/j.cell.2019.03.019

38. Nam, M.-H., Han, K.-S., Lee, J., Won, W., Koh, W., Bae, J. Y., Woo, J., Kim, J., Kwong, E., Choi, T.-Y., Chun, H., Lee, S. E., Kim, S.-B., Park, K. D., Choi, S.-Y., Bae, Y. C., & Lee, C. J. (2019). Activation of Astrocytic μ-Opioid Receptor Causes Conditioned Place Preference. Cell Reports, 28(5), 1154–1166.e5. 10.1016/j.celrep.2019.06.071

39. Navarrete, M., & Araque, A. (2010). Endocannabinoids Potentiate Synaptic Transmission through Stimulation of Astrocytes. Neuron, 68(1), 113–126. 10.1016/j.neuron.2010.08.043

40. Oe, Y., Wang, X., Patriarchi, T., Konno, A., Ozawa, K., Yahagi, K., Hirai, H., Tsuboi, T., Kitaguchi, T., Tian, L., McHugh, T. J., & Hirase, H. (2020). Distinct temporal integration of noradrenaline signaling by astrocytic second messengers during vigilance. Nature Communications, 11(1), 471. 10.1038/s41467-020-14378-x

41. Oh, S. W., Harris, J. A., Ng, L., Winslow, B., Cain, N., Mihalas, S., Wang, Q., Lau, C., Kuan, L., Henry, A. M., Mortrud, M. T., Ouellette, B., Nguyen, T. N., Sorensen, S. A., Slaughterbeck, C. R., Wakeman, W., Li, Y., Feng, D., Ho, A., … Zeng, H. (2014). A mesoscale connectome of the mouse brain. Nature, 508(7495), 207–214. 10.1038/nature13186

42. Owen, S. F., & Kreitzer, A. C. (2019). An open-source control system for in vivo fluorescence measurements from deep-brain structures. Journal of Neuroscience Methods, 311, 170–177. 10.1016/j.jneumeth.2018.10.022

43. Paukert, M., Agarwal, A., Cha, J., Doze, V. A., Kang, J. U., & Bergles, D. E. (2014). Norepinephrine Controls Astroglial Responsiveness to Local Circuit Activity. Neuron, 82(6), 1263–1270. 10.1016/j.neuron.2014.04.038

44. Paxinos, George., & Franklin, K. B. J. (2019). Paxinos and Franklin’s The mouse brain in stereotaxic coordinates. Academic Press, an imprint of Elsevier.

45. Pereira, M. J., Ayana, R., Holt, M. G., & Arckens, L. (2023). Chemogenetic manipulation of astrocyte activity at the synapse— a gateway to manage brain disease. Frontiers in Cell and Developmental Biology, 11. 10.3389/fcell.2023.1193130

46. Pin, J.-P., & Duvoisin, R. (1995). The metabotropic glutamate receptors: Structure and functions. Neuropharmacology, 34(1), 1–26. 10.1016/0028-3908(94)00129-G

47. Robin, L. M., Oliveira da Cruz, J. F., Langlais, V. C., Martin-Fernandez, M., Metna-Laurent, M., Busquets-Garcia, A., Bellocchio, L., Soria-Gomez, E., Papouin, T., Varilh, M., Sherwood, M. W., Belluomo, I., Balcells, G., Matias, I., Bosier, B., Drago, F., Van Eeckhaut, A., Smolders, I., Georges, F., … Marsicano, G. (2018). Astroglial CB1 Receptors Determine Synaptic D-Serine Availability to Enable Recognition Memory. Neuron, 98(5), 935–944.e5. 10.1016/j.neuron.2018.04.034

48. Rogan, M. T., Stäubli, U. V., & LeDoux, J. E. (1997). Fear conditioning induces associative long-term potentiation in the amygdala. Nature, 390(6660), 604–607. 10.1038/37601

49. Rosen, H., Stevens, R. C., Hanson, M., Roberts, E., & Oldstone, M. B. A. (2013). Sphingosine-1-Phosphate and Its Receptors: Structure, Signaling, and Influence. Annual Review of Biochemistry, 82(1), 637–662. 10.1146/annurev-biochem-062411-130916

50. Savtchouk, I., & Volterra, A. (2018). Gliotransmission: Beyond Black-and-White. The Journal of Neuroscience, 38(1), 14–25. 10.1523/JNEUROSCI.0017-17.2017

51. Shen, W., Li, Z., Tang, Y., Han, P., Zhu, F., Dong, J., Ma, T., Zhao, K., Zhang, X., Xie, Y., & Zeng, L. (2022). Somatostatin interneurons inhibit excitatory transmission mediated by astrocytic <scp> GABA B </scp> and presynaptic <scp> GABA B </scp> and adenosine <scp> A 1 </scp> receptors in the hippocampus. Journal of Neurochemistry, 163(4), 310–326. 10.1111/jnc.15662

52. Soares-Cunha, C., de Vasconcelos, N. A. P., Coimbra, B., Domingues, A. V., Silva, J. M., Loureiro-Campos, E., Gaspar, R., Sotiropoulos, I., Sousa, N., & Rodrigues, A. J. (2020). Nucleus accumbens medium spiny neurons subtypes signal both reward and aversion. Molecular Psychiatry, 25(12), 3241–3255. 10.1038/s41380-019-0484-3

53. Sugino, K., Clark, E., Schulmann, A., Shima, Y., Wang, L., Hunt, D. L., Hooks, B. M., Tränkner, D., Chandrashekar, J., Picard, S., Lemire, A. L., Spruston, N., Hantman, A. W., & Nelson, S. B. (2019). Mapping the transcriptional diversity of genetically and anatomically defined cell populations in the mouse brain. ELife, 8. 10.7554/eLife.38619

54. Sun, W., McConnell, E., Pare, J.-F., Xu, Q., Chen, M., Peng, W., Lovatt, D., Han, X., Smith, Y., & Nedergaard, M. (2013). Glutamate-Dependent Neuroglial Calcium Signaling Differs Between Young and Adult Brain. Science, 339(6116), 197–200. 10.1126/science.1226740

55. Suthard, R. L., Senne, R. A., Buzharsky, M. D., Pyo, A. Y., Dorst, K. E., Diep, A. H., Cole, R. H., & Ramirez, S. (2023). Basolateral Amygdala Astrocytes Are Engaged by the Acquisition and Expression of a Contextual Fear Memory. The Journal of Neuroscience, 43(27), 4997–5013. 10.1523/JNEUROSCI.1775-22.2023

56. Tasaka, G., Feigin, L., Maor, I., Groysman, M., DeNardo, L. A., Schiavo, J. K., Froemke, R. C., Luo, L., & Mizrahi, A. (2020). The Temporal Association Cortex Plays a Key Role in Auditory-Driven Maternal Plasticity. Neuron, 107(3), 566–579.e7. 10.1016/j.neuron.2020.05.004

57. Tasic, B., Menon, V., Nguyen, T. N., Kim, T. K., Jarsky, T., Yao, Z., Levi, B., Gray, L. T., Sorensen, S. A., Dolbeare, T., Bertagnolli, D., Goldy, J., Shapovalova, N., Parry, S., Lee, C., Smith, K., Bernard, A., Madisen, L., Sunkin, S. M., … Zeng, H. (2016). Adult mouse cortical cell taxonomy revealed by single cell transcriptomics. Nature Neuroscience, 19(2), 335–346. 10.1038/nn.4216

58. Taylor, C. R., Tse, V., Willoughby, D. D., Levesque, M., Vaidyanathan, T. V., Paz, J. T., & Poskanzer, K. E. (2025). Cortical astrocyte histamine-1-receptors regulate intracellular calcium and extracellular adenosine dynamics across sleep and wake. PLOS Biology, 23(10), e3003376. 10.1371/journal.pbio.3003376

59. Tovote, P., Fadok, J. P., & Lüthi, A. (2015). Neuronal circuits for fear and anxiety. Nature Reviews Neuroscience, 16(6), 317–331. 10.1038/nrn3945

60. Trumpp, N. M., Kliese, D., Hoenig, K., Haarmeier, T., & Kiefer, M. (2013). Losing the sound of concepts: Damage to auditory association cortex impairs the processing of sound-related concepts. Cortex, 49(2), 474–486. 10.1016/j.cortex.2012.02.002

61. Tyng, C. M., Amin, H. U., Saad, M. N. M., & Malik, A. S. (2017). The Influences of Emotion on Learning and Memory. Frontiers in Psychology, 8. 10.3389/fpsyg.2017.01454

62. Vaidyanathan, T. V, Collard, M., Yokoyama, S., Reitman, M. E., & Poskanzer, K. E. (2021). Cortical astrocytes independently regulate sleep depth and duration via separate GPCR pathways. ELife, 10. 10.7554/eLife.63329

63. Vita, N., Oury-Donat, F., Chalon, P., Guillemot, M., Kaghad, M., Bachy, A., Thurneyssen, O., Garcia, S., Poinot-Chazel, C., Casellas, P., Keane, P., Le Fur, G., Maffrand, J. P., Soubrie, P., Caput, D., & Ferrara, P. (1998). Neurotensin is an antagonist of the human neurotensin NT2 receptor expressed in Chinese hamster ovary cells. European Journal of Pharmacology, 360(2–3), 265–272. 10.1016/S0014-2999(98)00678-5

64. Wang, X., Lou, N., Xu, Q., Tian, G.-F., Peng, W. G., Han, X., Kang, J., Takano, T., & Nedergaard, M. (2006). Astrocytic Ca2+ signaling evoked by sensory stimulation in vivo. Nature Neuroscience, 9(6), 816–823. 10.1038/nn1703

65. Wu, Z., He, K., Chen, Y., Li, H., Pan, S., Li, B., Liu, T., Xi, F., Deng, F., Wang, H., Du, J., Jing, M., & Li, Y. (2022). A sensitive GRAB sensor for detecting extracellular ATP in vitro and in vivo. Neuron, 110(5), 770–782.e5. 10.1016/j.neuron.2021.11.027

66. Yao, Z., van Velthoven, C. T. J., Nguyen, T. N., Goldy, J., Sedeno-Cortes, A. E., Baftizadeh, F., Bertagnolli, D., Casper, T., Chiang, M., Crichton, K., Ding, S.-L., Fong, O., Garren, E., Glandon, A., Gouwens, N. W., Gray, J., Graybuck, L. T., Hawrylycz, M. J., Hirschstein, D., … Zeng, H. (2021). A taxonomy of transcriptomic cell types across the isocortex and hippocampal formation. Cell, 184(12), 3222–3241.e26. 10.1016/j.cell.2021.04.021

67. Yu, X., Taylor, A. M. W., Nagai, J., Golshani, P., Evans, C. J., Coppola, G., & Khakh, B. S. (2018). Reducing Astrocyte Calcium Signaling In Vivo Alters Striatal Microcircuits and Causes Repetitive Behavior. Neuron, 99(6), 1170–1187.e9. 10.1016/j.neuron.2018.08.015

68. Zingg, B., Hintiryan, H., Gou, L., Song, M. Y., Bay, M., Bienkowski, M. S., Foster, N. N., Yamashita, S., Bowman, I., Toga, A. W., & Dong, H.-W. (2014). Neural Networks of the Mouse Neocortex. Cell, 156(5), 1096–1111. 10.1016/j.cell.2014.02.023

